# Enhanced Population Control in Synthetic Bacterial Consortium by Interconnected Carbon Cross-Feeding

**DOI:** 10.1101/717926

**Authors:** Pauli S. Losoi, Ville P. Santala, Suvi M. Santala

## Abstract

Engineered microbial consortia can provide several advantages over monocultures in terms of utilization of mixed substrates, resistance to perturbations, and division of labor in complex tasks. However, maintaining stability, reproducibility, and control over population levels in variable conditions can be challenging in multi-species cultures. In our study, we modeled and constructed a synthetic symbiotic consortium with a genetically encoded carbon cross-feeding system. The system is based on strains of *Escherichia coli* and *Acinetobacter baylyi* ADP1, both engineered to be incapable of growing on glucose on their own. In a culture supplemented with glucose as the sole carbon source, growth of the two strains is afforded by the exchange of gluconate and acetate, resulting in inherent control over carbon availability and population balance. We investigated the system robustness in terms of stability and population control under different inoculum ratios, substrate concentrations, and cultivation scales, both experimentally and by modeling. To illustrate how the system might facilitate division of genetic circuits among synthetic microbial consortia, a green fluorescent protein sensitive to pH and a slowly-maturing red fluorescent protein were expressed in the consortium as measures of a circuit’s susceptibility to external and internal variability, respectively. The symbiotic consortium maintained stable and linear growth and circuit performance regardless of the initial substrate concentration or inoculum ratios. The developed cross-feeding system provides simple and reliable means for population control without expression of non-native elements or external inducer addition, being potentially exploitable in consortia applications involving precisely defined cell tasks or division of labor.

## Introduction

Synthetic biology enables the production of new-to-nature products with rationally designed multi-step pathways. Along with the increased complexity of the pathways and products, new challenges related to the functionality, stability, and over-burden of the cells have emerged. Sophisticated production platforms may require timed regulation and specialized synthesis patterns executed by multi-layer circuit designs (*1*). For all happening in one cell, the circuits are often prone to destabilization and loss of functionality, especially in up-scaled processes (*2*, *3*). Synthetic microbial consortia serve as an increasingly attractive platform for bioproduction due to their ability to address some of the challenges that are difficult to overcome with single strain cultures. Benefits of consortia include for example broader metabolic diversity (*4*), more efficient substrate utilization (*5–7*), tolerance to perturbations and metabolic imbalances (*8*), and possibility to distribute multi-task operations between different cells to alleviate metabolic burden (*9*). A number of applications ranging from bioremediation to drug production exploiting either natural or synthetic consortia have been recently reported (*10*). For example, genetically encoded task distribution between two microbial species has been employed in the production of diterpene chemotherapeutics, oxygenated taxanes (*11*). Neither of the strains was capable of producing oxygenated taxanes on its own, demonstrating the great potential of cocultures in the synthesis of complex products involving multiple reaction steps.

Despite the several advantages of coculture systems, challenges related to population control and complex (and potentially counterproductive) interactions between the different strains exist. In addition, optimizing the growth conditions for all subpopulations is more complicated than for single-strain cultures. Genetic circuits exploiting for example cell-to-cell signaling can be applied to establish spatiotemporal control over the functions of different cells (*12–14*). However, it is desirable to engineer the cells for symbiotic relationship in order to improve stability and to control population ratios, competition for nutrients, and disadvantageous interactions (*15*). One approach is to provide separate carbon sources for consortium members (*5*, *16*), sometimes allowing one strain to grow only on side-products of another strain (*11, 17*). Examples of genetically encoded nutrient cross-feeding based on amino acid or essential metabolite exchange have also been reported (*18–20*).

Modeling and predicting consortium behavior and performance is also more complicated than for monocultures. Unstructured kinetic models are relatively straightforward to assemble and apply (*4*, *18*, *19*, *21*), but their mechanistic formulation demands prior knowledge of the consortium at hand. Genome-scale models have more predictive potential, but their correct application in community flux balance analysis (FBA) methods requires careful consideration of both the solution procedure and the variables and quantities involved. Most multi-strain FBAs do not decouple the community members from each other, but allow for direct exchange between strains during an optimization step (*19, 22*), which may result in extensive and non-realistic cross-feeding as some community members waste resources to support the rest of the community (*23*). In order to avoid such “altruism” without employing non-linear objective functions (*24*), the strains need to be optimized for separately, in a decoupled manner (*25*). However, this complicates accounting for community structure without resorting to dynamic FBAs.

For a multi-step production pathway, balanced production of intermediates as well as having a sufficiently long and stable production window is important (*26*). When a pathway is split between several hosts (*11, 16*), sustaining the balance becomes even more crucial, but also more challenging. Thus, strict control over the growth of subpopulations is required. Some cocultures of two strains have involved a mixture of mutualistic and synergistic population dynamics, in which only one of the strains is unconditionally dependent on the interactions, whereas the other at most benefits from coexistence (*4*, *11*, *17*). In this type of oneway mutualism, the control over subpopulations is somewhat loose, and may not necessarily endure in longer cultivations or with varying substrate concentrations. On the other hand, mutually obligate cross-feeding of amino acids (*20*) or other essential metabolites (*18, 19*) may not be in balance in terms of quantity and quality and might interfere with the (over)expression of the production pathway enzymes. In addition, the strains are in a sense competing over the exchanged essential metabolites and also for the same carbon substrate, which could result in differential growth rates or reduced production rates.

In this study, our aim was to model and construct a synthetic consortium with an interconnected cross-feeding system that is solely based on the cooperative utilization of a single carbon source. We employed as hosts *Escherichia coli* K-12 and the soil bacterium *Acinetobacter baylyi* ADP1, which has been previously used both in synthetic microbial consortia (*17*, *27*, *28*) and in production of industrially relevant compounds from various substrates (*29–31*). In order to implement the cross-feeding between *E. coli* and *A. baylyi* ADP1, both strains were engineered to be incapable of glucose utilization alone. In minimal salts medium supplemented with glucose, *A. baylyi* ADP1 oxidizes glucose to gluconate, which is then transported into the cells, but the ∆*gntT* mutant (high-affinity gluconate permease gene deleted) is incapable of importing the gluconate. *E. coli* ∆*ptsI* (PTS enzyme I gene deleted), in turn, cannot utilize glucose, but can grow on gluconate made available by *A. baylyi* ADP1 ∆*gntT*. While utilizing gluconate, *E. coli* ∆*ptsI* produces acetate which serves as the carbon source for *A. baylyi* ADP1 ∆*gntT*. Thus, the availability of carbon source is dependent on both strains as illustrated in Figure 1, resulting in inherent growth control. We investigated the consortium behavior and robustness in terms of stability and population control by modeling with a combination of FBA and unstructured kinetics and by performing experiments under different inoculum ratios, substrate concentrations, and cultivation scales. As a demonstration of how the population control provided by the interconnected carbon cross-feeding system might apply to division of labor among a synthetic microbial consortium, the superfolder green fluorescent protein sfGFP (*32*) and the monomeric red fluorescent protein mScarlet (*33*) were used as a surrogate of a genetic circuit distributed in the consortium. The sfGFP’s pH-sensitivity (*34*) and the mScarlet’s long maturation time (*35*) corresponded then to the circuit’s susceptibility to external conditions and internal variability (*2*), respectively.

**Figure 1:**
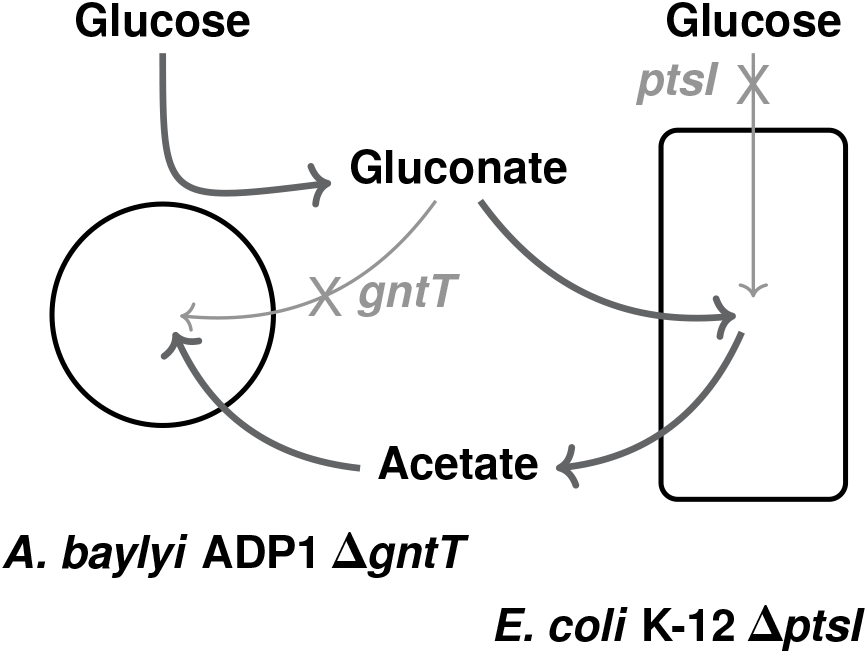
Carbon flow in the synthetic consortium of engineered *Escherichia coli* (Ec) and *Acinetobacter baylyi* ADP1 (Ab) based on cooperative utilization of glucose. Ab oxidizes glucose to gluconate, but the ∆*gntT* (high-affinity gluconate permease) mutant is incapable of importing gluconate. The Ec∆*ptsI* (PTS enzyme I) mutant cannot utilize glucose but is capable of growing on the gluconate made available by Ab∆*gntT*. Upon growth on gluconate Ec∆*ptsI* produces acetate, which in turn serves as carbon source for Ab∆*gntT*.

## Results and Discussion

To provide support for the hypothetized growth-enabling carbon cross-feeding between the glucose negative strains, *A. baylyi* ADP1 (Ab) ∆*gntT* and *E. coli* (Ec) ∆*ptsI*, FBAs were carried out. Adopting the decoupled iterative scheme (*25*) with a slight modification, FBAs predicted that a consortium of Ab∆*gntT* and Ec∆*ptsI* should grow on glucose. According to the FBAs, growth was enabled by conversion of glucose to gluconate by Ab, growth on gluconate by Ec, and growth of Ab on acetate excreted by Ec (Figure 1). The results were sensitive to oxygen availability (Supplementary Section S1), as metabolite excretion is dependent of allowed oxygen input in FBAs (*25*).

Having established that carbon cross-feeding should allow growth of the knock-out strains in coculture, the Ab∆*gntT*:Ec∆*ptsI* consortium’s behaviour was investigated from a theoretical perspective. Provided that the total biomass concentration *X*_T_ = *X*_1_ + *X*_2_ (mass/volume) of a two-strain culture is increasing or decreasing, the change in the strain *i*’s biomass proportion *X_i_/X*_T_ (mass/mass) with respect to time *t* can be written as (Supplementary Section S2)

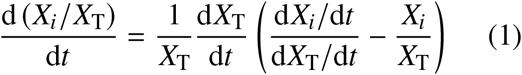

This formulation resembles mass or heat transfer equations towards an equilibrium state, in which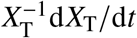 represents the time scale (1/time) and the difference in parentheses the driving force of transfer towards equilibrium. As the subtracted term in the driving force is the strain *i*’s proportion of total biomass, *X_i_/X*_T_, the term (d*X_i_*/d*t*)/(d*X*_T_/d*t*) can be interpreted as the equilibrium state the consortium is adapting to. Identifying whether this so-called equilibrium state is variable or constant allows predicting whether the relative population levels are volatile or stable, respectively.

Considering an unstructured kinetic model of the Ab∆*gntT*:Ec∆*ptsI* consortium in batch culture with growth according to the Monod equation, conversion of glucose to gluconate (N) by Ab, consumption of gluconate and excretion of acetate (A) by Ec, and consumption of acetate by Ab, the equilibrium proportion of Ab simplifies to (Supplementary Sections S2.1 and S2.2)

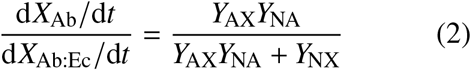

given that the concentration of acetate is constant (d*A*/d*t* = 0). In eq 2, *Y*_AX_ is the yield of Ab biomass on acetate, *Y*_NX_ is the yield of Ec biomass on gluconate, and *Y*_NA_ is the yield of acetate on gluconate (mass/mass). Numerical integrations of the ordinary differential equation (ODE) system consisting of concentrations of Ab and Ec biomasses, glucose, gluconate, and acetate resulted invariably in acetate reaching a steady state of constant concentration after an initial accumulation period (Supplementary Section S2.3). Equation 2 then effectively states that the optimal composition of Ab∆*gntT*:Ec∆*ptsI* is determined solely by how much acetate is produced from gluconate by Ec∆*ptsI* (*Y*_NA_) and how much biomass can be formed from acetate by Ab∆*gntT* (*Y*_AX_) and from gluconate by Ec∆*ptsI* (*Y*_NX_). If the yield coefficients are assumed to be constants, eq 2 implies tight population control towards uniquely defined and stable optimum strain ratios.

Similar expressions of equilibrium strain proportions were also derived for three other consortia of *A. baylyi* ADP1 and *E. coli* by pairing both wild-type and knock-out strains. The resulting consortia Ab:Ec, Ab∆*gntT*:Ec, and Ab:Ec∆*ptsI* corresponded to competition, commensialism, and cooperator-cheater dynamics, respectively (Supplementary Section S2.2). After solving the associated ODE systems numerically, it was concluded that none of these consortia without the two-way coupling had a uniquely defined and stable equilibrium state in the same sense as Ab∆*gntT*:Ec∆*ptsI*. The equilibrium states were found to be dependent of either acetate fluctuations, as d*A*/d*t* ≈ 0 did not apply, or the ever changing biomass concentrations (Supplementary Sections S2.2 and S2.3). As the equilibrium states were not constant but were changing over time, the other consortia were under looser control. These simple kinetic analyses highlighted how the interconnected carbon cross-feeding system present in Ab∆*gntT*:Ec∆*ptsI* should enhance population control by defining a constant strain ratio optimum. As a consequence, distributed expression of circuits or pathways can be expected to be more balanced in Ab∆*gntT*:Ec∆*ptsI*.

In order to experimentally study the consortia and confirm the theoretical predictions presented above, *A. baylyi* ADP1 and *E. coli* were genetically engineered to constitutively express mScarlet and sfGFP, respectively. The genes encoding mScarlet and sfGFP were integrated into genomes to avoid unpredictable plasmid copy number effects (*36*) and to more accurately represent the performance of each strain in the consortium. A gene cassette (*37*) carrying sfGFP under the constitutive promoter BBa_J23100 and a gentamicin resistance gene was introduced to wild-type Ec and knock-out Ec∆*ptsI* genomes by *ϕ*80 phage integrase (*38*), yielding the fluorescent strains designated as Eg and Ekg, respectively. For Ab and Ab∆*gntT*, sfGFP was replaced by mScarlet, and a gene cassette (*39*) containing appropriate flanking regions and a chloramphenicol resistance gene was utilized to facilitate genomic integration to the neutral (*39*) locus ACIAD3381 (*poxB*, pyruvate dehydrogenase) by natural transformation. The resulting fluorescent Ab and Ab∆*gntT* strains were designated as Ar and Akr. Table 1 summarizes the strains constructed and utilized in this work.

**Table 1:**
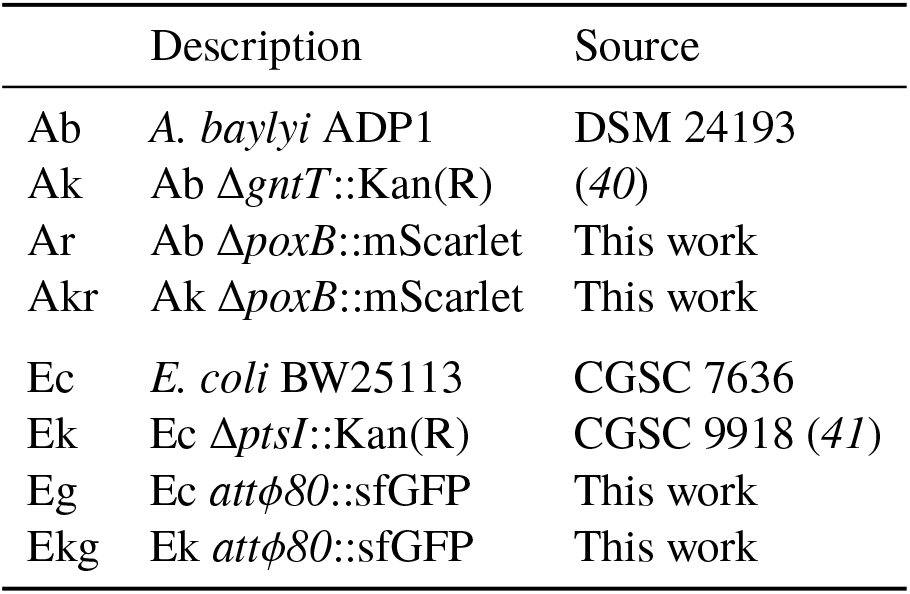
The constructed and utilized strains. The kanamycin resistant Kan(R) glucose knock-out strains are designated with k, sfGFP-carrying strains with g, and mScarlet-carrying strains with r. The mScarlet and sfGFP cassettes carried also resistance genes for chloramphenicol and gentamicin, respectively. DSM stands for Deutsche Sammlung von Mikroorganismen und Zellkulturen and CGSC for Yale University Coli Genetic Stock Center.

The 8 strains were then cultivated for 45–47 h in small scale (200 µL) at 30 °C in a minimal medium with 50 mm glucose as the sole carbon source. All strains were cultured in isolation and in all 16 possible Ab:Ec-pairs. Akin to previous studies involving obligately cross-feeding consortia (*18–20*), none of the knock-out strains could grow in isolation, but each consortium, and remarkably also the double knock-out ones, accumulated a considerable optical density at 600 nm (OD_600_). However, all consortia with two knock-out strains (Ak or Akr with Ek or Ekg) also grew slower (Supplementary Figure S5) than the other consortia with one or no knock-out strains (*19*). This can be attributed to both Ec∆*ptsI* and Ab∆*gntT* presumably growing only on substrates derived from glucose, namely gluconate and acetate, respectively, which inevitably were less available than glucose initially. Fluorescence intensities corresponding to emission maxima of sfGFP and mScarlet also grew considerably, but only if a strain with the corresponding fluorescent protein gene was present in the culture (Supplementary Figure S6). The steady rise of not only OD_600_, but also of both sfGFP and mScarlet fluorescences when applicable therefore indicated that indeed both knock-out strains grew in the Ab∆*gntT*:Ec∆*ptsI* consortia.

The developed cross-feeding system scaled-up successfully as well, as an 18 h cultivation of Akr:Ekg at the same conditions in a 0.5 L aerated bioreactor resulted likewise in growth of both OD_600_ and fluorescences. High-performance liquid chromatograph analyses (Supplementary Section S6) of the cultivation further confirmed the carbon flow scheme suggested in Figure 1 and supported by FBAs. The cross-feeding not only enabled growth of knock-out strains, but reduced side product (acetate) accumulation, which translated to an overall enhanced carbon utilization often associated with cocultures (*5*, *11*, *17*). On the other hand, the slowness of growth afforded by the two-way crossfeeding is disadvantageous from a bioprocessing perspective, as it could lead to prohibitively low overall productivities if the system were to be applied in production of bulk chemicals (*42*). Identifying the limiting intermediate, likely acetate in this case (Supplementary Section S6), and increasing its rate of production (*11*) could elevate the overall growth rate. Another challenge is that the present implementation is based on pure glucose as the carbon source. This could complicate the system’s application at large scale, as cost considerations make heterogeneous feed mixtures preferable in industry (*43*).

Having confirmed the growth of and carbon flow within Ab∆*gntT*:Ec∆*ptsI*, its stability was then experimentally tested with respect to time and initial substrate concentration by monitoring fluoresence of the mScarlet:sfGFP model circuit embedded in Akr:Ekg. The small-scale experiments were repeated under initial glucose concentrations of 50 mm, 100 mm, and 200 mm, and the OD_600_ and fluorescences measured from Akr:Ekg and Ar:Eg (Ab∆*gntT*:Ec∆*ptsI* and Ab:Ec with mScarlet and sfGFP) are shown in Figure 2. The initial glucose concentration had no discernible effect on growth or fluorescence of either Akr:Ekg or Ar:Eg. Regardless of initial concentration of glucose, Ar:Eg followed a typical batch growth pattern of lag, exponential, and stationary phases, whereas Akr:Ekg grew in a much more steady and linear fashion. The same trends applied also to sfGFP and mScarlet fluorescences. As the unstructured kinetics implied by predicting a fixed optimum of strain ratios in Ab∆*gntT*:Ec∆*ptsI*, Akr:Ekg maintained the balance between mScarlet and sfGFP. Averaged over the three initial glucose concentrations, the ratio of mScarlet to sfGFP varied by a factor of 3.3 1.3 (max min) in Akr:Ekg, whereas in Ar:Eg the factor was over 5-fold larger (18.4 ± 7.4) as mScarlet reached maximum activity several hours after sfGFP. The uncertainty derives from mScarlet and sfGFP sample standard deviations propagating error to mScarlet-sfGFP and max-min ratios and to averaging max min over the three experiments containing six biological replicates each. In terms of balanced expression required by multi-layer circuits and multi-step production pathways (*11*–*14*, *16*, *26*), Akr:Ekg was superior to the wild-type Ar:Eg consortium.

**Figure 2:**
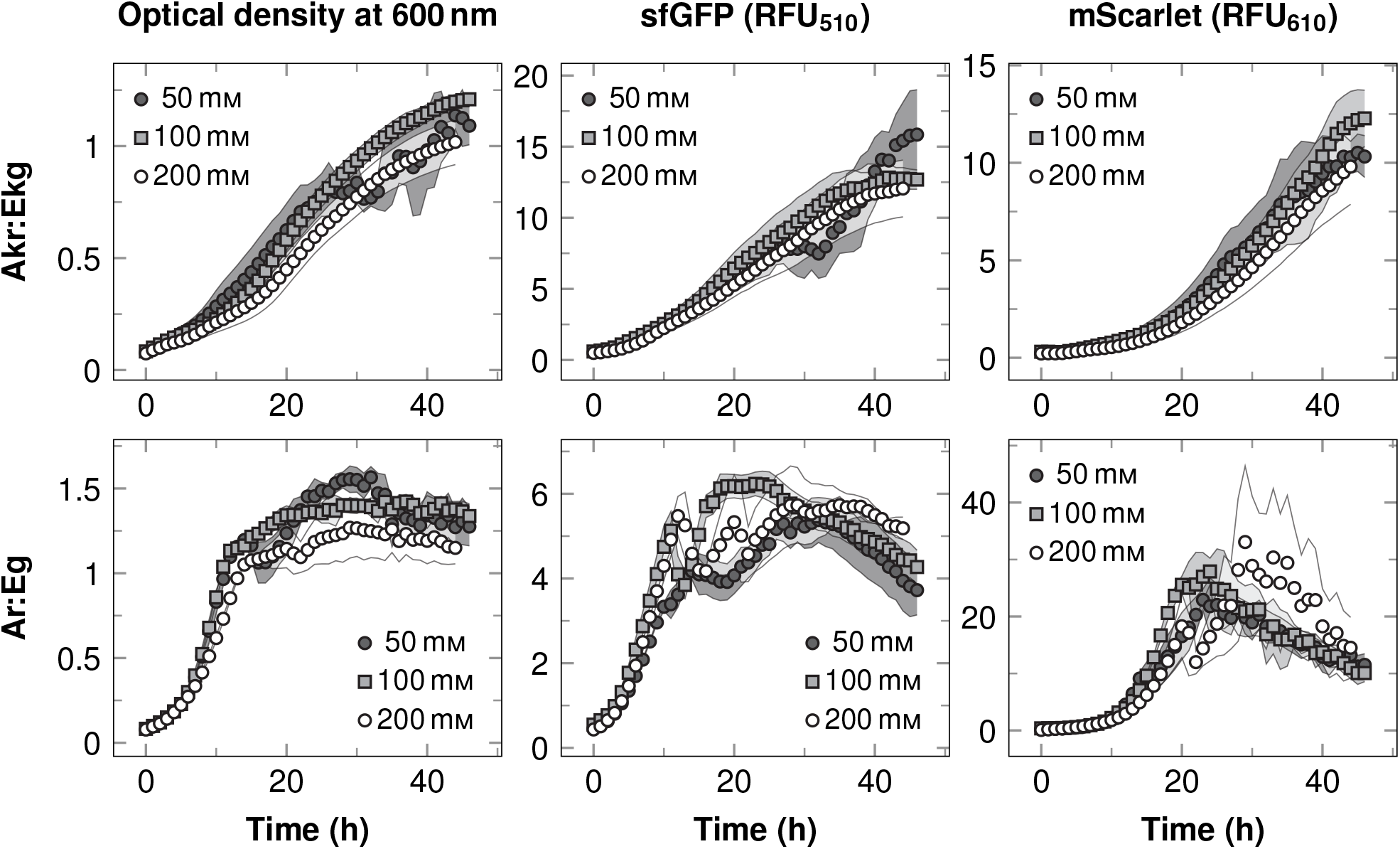
Effect of initial glucose concentration on consortia of *A. baylyi* ADP1 (Ab) and *E. coli* (Ec) expressing mScarlet and sfGFP, respectively. Ab∆*gntT*:Ec∆*ptsI* (Akr:Ekg, top row) and Ab:Ec (Ar:Eg, bottom row) were cultivated under different initial glucose concentrations in a defined medium at 30 ◦C. Fluorescence intensities at 510 nm and 610 nm (sfGFP and mScarlet, respectively) are reported in the relative fluorescence units (RFU) of the microwell plate reader divided by 1000. Each line stands for an initial glucose concentration, and the data are shown as means of six biological replicates with shaded areas representing sample standard deviations. At some points the error bands are smaller than the markers. For clarity, the data are presented hourly.

It was then tested whether the presented carbon flow in Ab∆*gntT*:Ec∆*ptsI* would stabilize growth and distributed expression by exerting control also over initial population composition as predicted by the kinetic model. The Akr:Ekg consortium and Ar:Eg for reference were cultivated with initial Ec to Ab ratios of 1:1, 2:1, 5:1, and 9:1 in terms of OD_600_ using 100 mm glucose as the carbon source at otherwise same conditions. The courses of these cultivations are shown in Figure 3, and to further emphasize the difference between Akr:Ekg and Ar:Eg, the ratio of mScarlet to sfGFP is depicted in Figure 4. The variation in inoculum composition caused practically no variation at all in Akr:Ekg performance: OD_600_ and both fluorescences rose as almost linear functions of time with nearly identical slopes. With higher Ab inoculum proportions the ratio of mScarlet to sfGFP fluoresence was proportionally higher at first, but Akr:Ekg stabilized the ratio remarkably well to similar low-sloped linear functions of time. Ar:Eg, however, displayed imbalances in mScarlet and sfGFP activities similarly to the earlier experiments with varying initial substrate concentrations. Additionally, a sharp decline in sfGFP activity was observed at 15 h with higher initial Ec proportions. This was likely caused by Ec acidifying the culture (*17*), which in turn would have reduced sfGFP fluorescence (*34*): pH declined clearly in larger-scale cultivations of Ar:Eg (with equal proportions of Ab and Ec initially) and Ec in isolation (Supplementary Figure S7). In contrast to Ar:Eg, the long and stable production window demonstrated by Akr:Ekg for up to 40 h underlined how the two-way cross-feeding system stabilized the distributed circuit also against external factors (modeled by sfGFP’s pH sensitivity) and internal variability (represented by mScarlet’s long maturation time).

**Figure 3:**
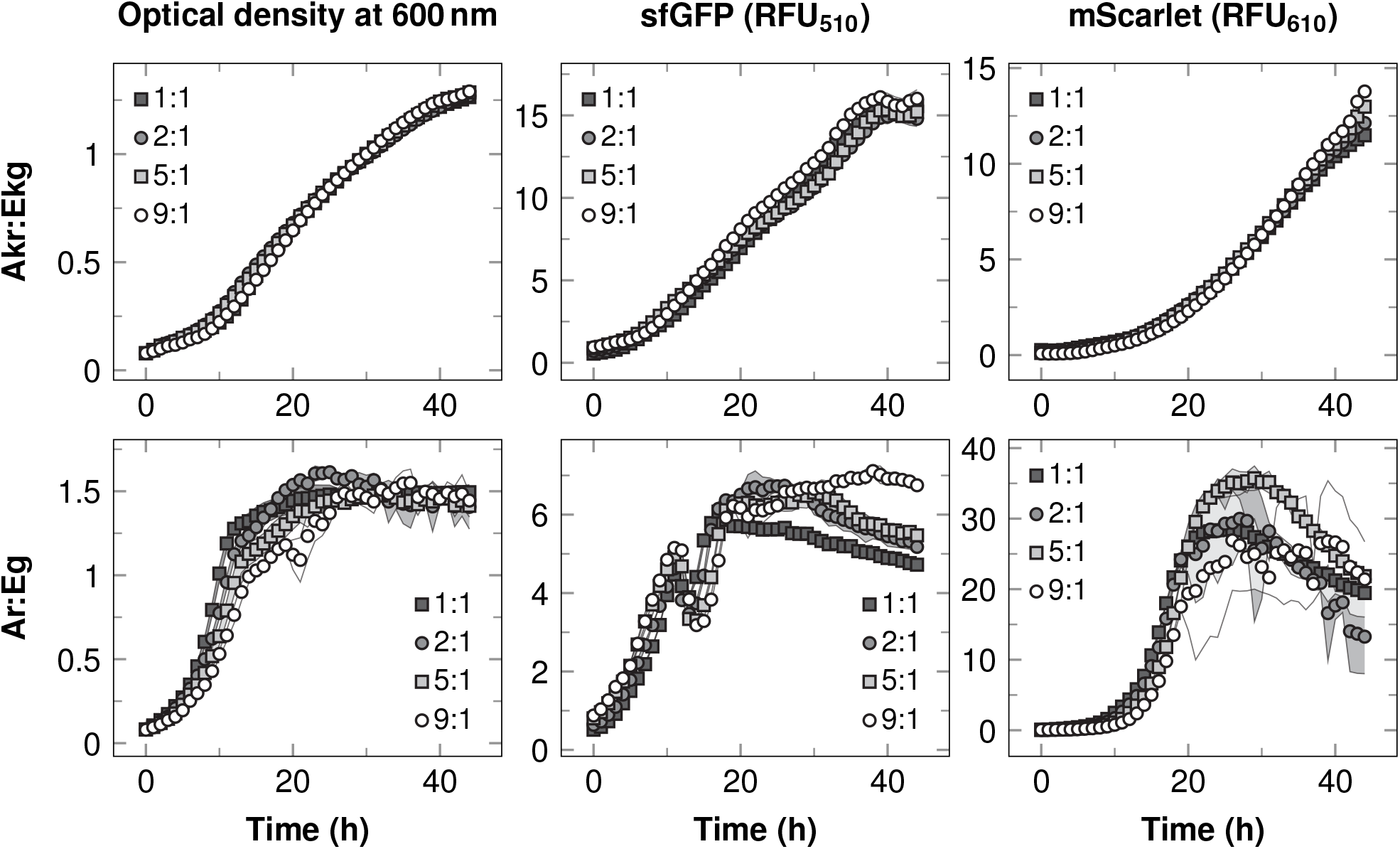
Effect of inoculation ratios on consortia of *E. coli* (Ec) and *A. baylyi* ADP1 (Ab) expressing sfGFP and mScarlet, respectively. The wild-type Ar:Eg (Ab:Ec, bottom row) and knock-out Akr:Ekg (Ab∆*gntT*:Ec∆*ptsI*, top row) pairs were cultivated with initial Ec to Ab ratios of 1:1, 2:1, 5:1, and 9:1 in terms of OD_600_ in a defined medium with 100 mm glucose at 30 °C. RFUs at 510 nm and 610 nm (sfGFP and mScarlet, respectively) stand for the microwell plate reader’s relative fluorescence units divided by 1000. The data are presented as means of three biological replicates with shaded areas representing sample standard deviations. For Akr:Ekg the error bands are smaller than the markers. The data are shown hourly for clarity.

**Figure 4:**
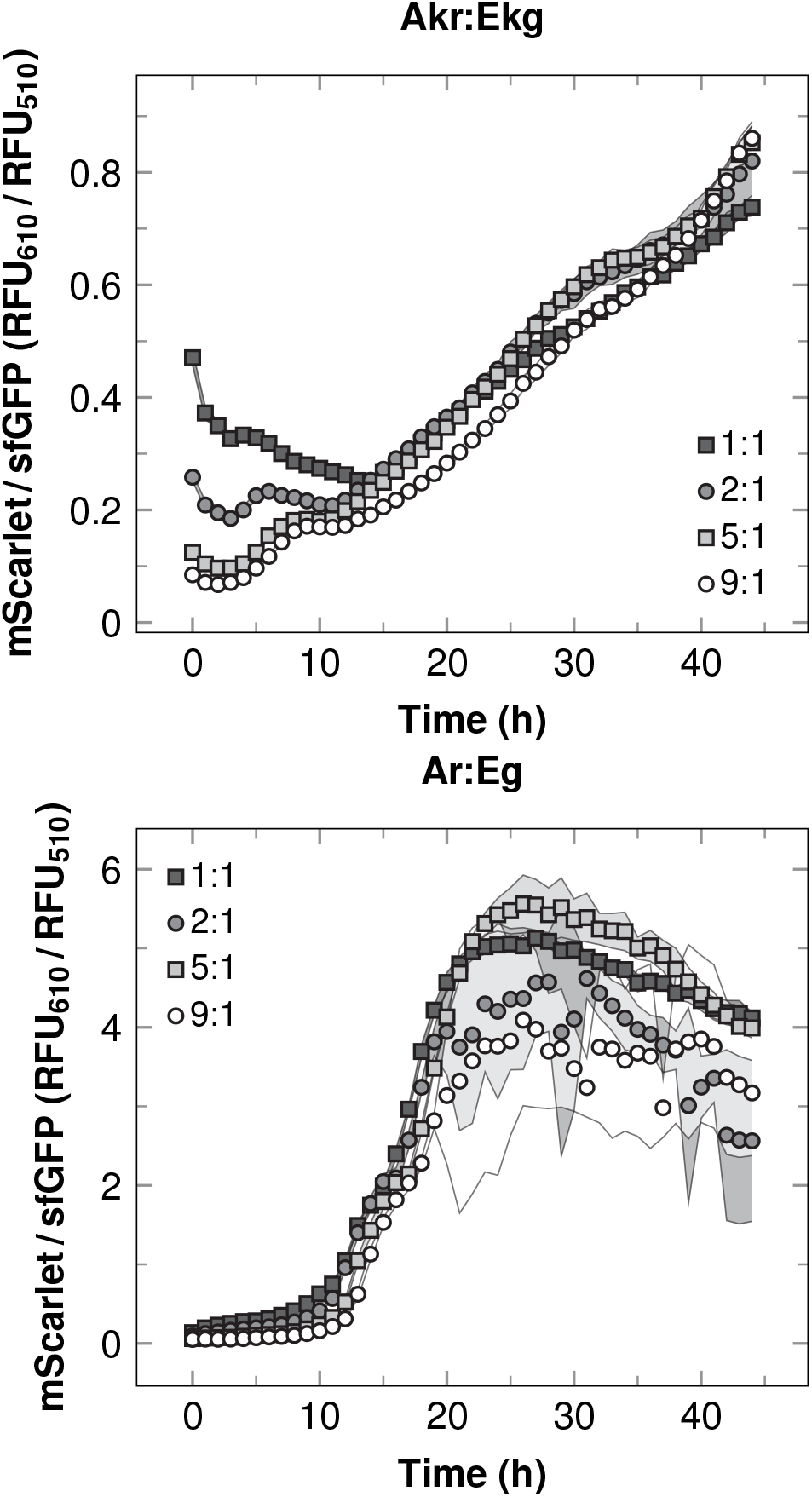
Effect of consortium inoculation ratios on the balance between mScarlet and sfGFP expressed by *A. baylyi* ADP1 (Ab) and *E. coli* (Ec), respectively. Cultivations of Akr:Ekg (Ab∆*gntT*:Ec∆*ptsI*, top) and Ar:Eg (Ab:Ec, bottom) were performed with Ec to Ab inoculation ratios of 1:1, 2:1, 5:1, and 9:1 in terms of OD_600_ at 30 ◦C in a defined medium with 100 mm glucose. RFUs at 510 nm and 610 nm are the relative fluorescence units of the microwell plate reader divided by 1000. Three biological replicates were cultivated, and the presented data are ratios of mScarlet means to sfGFP means. Shaded areas correspond to sample standard deviations. For Akr:Ekg the error bands are mostly smaller than the markers. The data are presented hourly for clarity.

Unlike the cross-feeding system presented in this work, where neither of the strains grows directly on the provided substrate, many cross-feeding concepts have involved a one-way coupling of carbon flow, where one of the strains grows on the supplied substrate and the other one grows on products of the initially growing strain (*4*, *11*, *17*, *28*). Theoretically such one-way carbon flow can result in constant and unique strain ratios, analogous to eq 2, and published performances of such consortia also point at stable functioning. However, stability is not as generally guaranteed as in two-way cross-feeding, because growth and metabolism of one strain is not constrained by the other (Supplementary Sections S2.2 and S2.3). Other two-way cross-feeding concepts have been based on exchange of biosynthetically costly metabolites (*18–20*) required and utilized by all consortium members simultaneously. Exchange of essential metabolites carries a cost to the contributing strains, however, and such metabolites are likely not as readily exchanged as gluconate and acetate, which are costless (*25*) side-products not utilized by the contributing Ab and Ec strains themselves (*19*) in Ab∆*gntT*:Ec∆*ptsI*.

The presented carbon cross-feeding system has therefore several advantages compared to other forms of population control: it does not require expression of non-native genes as it is based entirely on native host metabolism, it imposes predictable and robust optimal strain ratios, it does not demand the strains to donate essential metabolites required by themselves, and it provides easily modelable and optimizable, almost linear growth and expression patterns. The system was functional even across cultivation scales, and it should be applicable in balancing distributed expression of circuits (*9*) or pathways (*6*, *11*, *16*) without separate feeds (*5, 16*) or adjustment of inoculation ratios (*7, 16*). The cross-feeding system should also be less susceptible to fluctuating conditions (*2*) and deleterious mutations (*3*) than systems based on external inducers (*2*, *6*, *26*) or complex genetic circuitry (*1*, *2*, *9*, *12*–*14*), as it is implemented by only two simple gene deletions. Conversely, the developed system is host-specific and not necessarily directly transferable to other species, but introducing non-native genes in implementing the cross-feeding might enable its generalization. Based on the presented theoretical aspects and experimental results, it is concluded that interconnected carbon cross-feeding provides simple yet reliable population control, and consequently an attractive platform for balanced and robust expression of distributed genetic circuits and production pathways.

## Methods

### Strain construction

The wild type strains designated as Ab and Ec were *A. baylyi* ADP1 (DSM 24193) and *E. coli* K-12 BW25113 (CGSC 7636), respectively. The knockout strain Ab∆*gntT* (ACIAD0544 high-affinity gluconate permease gene deleted) (*40*) was a kind gift from Veronique de Berardinis (Genoscope, France) and Ec∆*ptsI* (PTS enzyme I gene deleted) (*41*) was obtained from Yale University Coli Genetic Stock Center (CGSC 9918). In both knock-out strains the deleted gene is replaced by a kanamycin resistance gene. All the fluorescent strains Ar, Akr, Eg, and Ekg constructed in this work had a genomically integrated fluorescent protein expression cassette along with a chloramphenicol (Ab) or gentamicin (Ec) resistance gene for selection.

Genomic integrations of the sfGFP expression cassette into Ec and Ec∆*ptsI* were performed with conditional-replication, integration, and modular (CRIM) plasmids as described in (*38*). The CRIM plasmids Burden Monitor phi80 version (Addgene plasmid #66074) and pAH123 (Addgene plasmid #66077) were gifts from Tom Ellis (*37*). Supplementary Section S4 contains additional details.

The mScarlet cassette was similar to the sfGFP cassette (*37*) in design (Supplementary Section S3), and it was ordered from GenScript (New Jersey, USA). For integration into *A. baylyi* ADP1 genome, the mScarlet cassette was inserted within a gene cassette (*39*) overwriting the neutral ACIAD3381- locus (*poxB*, pyruvate dehydrogenase). The cassette was integrated into Ab and Ab∆*gntT* by natural transformation in solid phase by pipetting 300 ng of the transforming DNA directly on single colonies on LA plates (15 g/L agar, 10 g/L tryptone, 5 g/L yeast extract, 1 g/L NaCl, and 10 g/L glucose). The plates were incubated overnight in room temperature, and the colonies were replated on other LA plates with 25 mg/L chloramphenicol for selection.

### Small-scale cultivations

Small-scale cultivations were performed with 200 µl culture volumes on 96-well plates in a mineral salts medium (*44*) (Supplementary Section S5) with 50 mm, 100 mm, or 200 mm glucose as the carbon source. The plates were incubated in a Spark multi-mode microplate reader (Tecan, Switzerland) with temperature control set to 30 °C. Optical density at 600 nm, and fluorescence intensities with excitation-emission filter-pairs of 485-510 nm (sfGFP) and 580-610 nm (mScarlet) were recorded every 30 minutes. Gain was set to a fixed value of 50 for the fluoresence measurements. Plates were shaken in between measurements. Precultivations were performed in the same medium with 10 mm Na-acetate, 10 mm Na-gluconate, and 10 mm glucose as the carbon sources at 30 °C with 300 RPM shaking. The targeted initial OD_600_ for the microwell plates was 0.1.

### Numerical work

Data analyses and ODE integrations were performed using the Python programming language (http://www.python.org) along with scipy (*45*), numpy (*46*), and pandas (*47*) packages. The cobrapy package (*48*) was utilized for FBAs. Sample standard deviations *s _f_* of functions *f* with variables *x_i_* were calculated including error propagation with zero covariance: 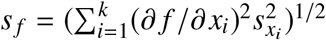.

FBAs were conducted according to the iterative scheme from (*25*), but with an additional step prepended to each iteration: first the glucose importing exchange reaction was maximized for and the maximized value was set as a fixed bound for the subsequent growth rate maximizing optimization step. After growth reaction maximization, an additional minimization of the sum of all gene-associated flux was carried out as in (*25*), but given also the constraint of maximized glucose import. Contrary to the method presented in (*25*), secreted metabolites were incorporated into computational medium in subsequent iteration regardless of growth status. The utilized genome-scale models of *A. baylyi* ADP1 and *E. coli* were iAbaylyiv4 (*49*) and iAF1260 (*50*), respectively. Computational medium was made to resemble the defined medium used in experiments. Supplementary Section S1 contains additional details.

## Supporting Information Available

Additional results, discussion, and methods for flux balance analyses, kinetic modeling, strain and plasmid construction, and experiments.

## Author Contributions

P.S.L., V.P.S., and S.M.S. designed the study. P.S.L. performed modeling, strain construction, and larger-scale experiments. V.P.S. and S.M.S. performed small-scale experiments and supervised the study. All authors participated in preparing the manuscript. All authors approved the final manuscript.

## Notes

The authors declare no competing financial interest.

## Acknowledgements

The authors thank Tapio Lehtinen for his comments on the manuscript. Funding was provided by Academy of Finland (grant numbers 310135 and 310188) and Tampere University of Technology Graduate School (P.S.L.).

