## Supplementary Text for "Enhanced Population Control in Synthetic Bacterial Consortium by Interconnected Carbon Cross-Feeding"

**S3 mScarlet Gene Cassette Construction** **15**

**S4 sfGFP Integrations** **17**

**S5 Microwell Cultivations** **18**

**S6 Bioreactor Cultivations** **18**

**Supplementary References** **23**

### S1 Flux Balance Analyses

This section describes how the carbon flow of the double knock-out Ab$\Delta$*gntT*:Ec$\Delta$*ptsI* consortium was predicted using flux balance analyses (FBAs) with an iterative scheme [1]. Table S1 shows the iteration progress and the appearance of gluconate and acetate. In the double knock-out Ab$\Delta$*gntT*:Ec$\Delta$*ptsI* consortium, neither *Acinetobacter baylyi* ADP1 $\Delta$*gntT* (Ab$\Delta$*gntT*) nor *Escherichia coli* $\Delta$*ptsI* (Ec$\Delta$*ptsI*) was capable of growing in the first iteration. Ak converted glucose to gluconate, however. Given the gluconate made available by *A. baylyi* ADP1, *E. coli* was capable of growing and produced acetate as a consequence (when oxygen input was less than or equal to 12.5 mmol $g_{DW}^{-1}$ $h^{-1}$). Growth of *A. baylyi* ADP1 is then observed in the third step, as acetate has been made available by *E. coli*. As for the wild-type Ab:Ec consortium, both *A. baylyi* ADP1 and *E. coli* grew independetly.

Despite all its merits, the growth rate maximization commonly employed in FBAs has a drawback: it implicitly assumes a perfectly adapted population, which in the case of *E. coli* resulted in no acetate production given a sufficient oxygen input. Not all of acetate produced by *E. coli* can be explained by fermentative metabolism alone under glucose excess [2–4], and to compensate for this defect the FBAs were conducted at different oxygen availabilities.

Table S1: Growth statuses and secreted metabolites in the iterative [1] FBAs. Growth of a strain in a particular iteration step (rows) with a given oxygen input (columns) is indicated by boldface font of the listed metabolites. For example, Ak (*A. baylyi* ADP1 Δ*gntT*) grew in Ak:Ek at third iteration with oxygen inputs of 10 mmol $g_{DW}^{-1}$ $h^{-1}$ and 12.5 mmol $g_{DW}^{-1}$ $h^{-1}$ but not with 15 mmol $g_{DW}^{-1}$ $h^{-1}$, whereas Ek (*E. coli* Δ*ptsI*) grew with all oxygen inputs. Both wild-type strains Ab (*A. baylyi* ADP1) and Ec (*E. coli*) grew in Ab:Ec during all iterations as neither was dependent of the other. Formate was not utilized by any of the consortium members even though it was secreted in the FBA simulations. Abbreviations: Glcn gluconate, Ace acetate, For formate.

| Iteration | Strain | 10 $mmol_{O_2}$ $g_{DW}^{-1}$ $h^{-1}$ | 12.5 $mmol_{O_2}$ $g_{DW}^{-1}$ $h^{-1}$ | 15 $mmol_{O_2}$ $g_{DW}^{-1}$ $h^{-1}$ |
|---|---|---|---|---|
| Ak:Ek | | | | |
| 1 | Ak | Glcn | Glcn | Glcn |
| | Ek | | | |
| 2 | Ak | Glcn | Glcn | Glcn |
| | Ek | **$H_2O$, $CO_2$, Ace, $H^+$** | **$H_2O$, $CO_2$, Ace** | **$H_2O$, $CO_2$** |
| 3 | Ak | **$H_2O$, $CO_2$, Glcn, For** | **$H_2O$, $CO_2$, Glcn, For** | Glcn |
| | Ek | **$H_2O$, $CO_2$, Ace, $H^+$** | **$H_2O$, $CO_2$, Ace** | **$H_2O$, $CO_2$** |
| 4 | Ak | **$H_2O$, $CO_2$, Glcn, For** | **$H_2O$, $CO_2$, Glcn, For** | |
| | Ek | **$H_2O$, $CO_2$, Ace, $H^+$** | **$H_2O$, $CO_2$, Ace** | |
| Ab:Ec | | | | |
| 1 | Ab | **$H_2O$, $CO_2$, Glcn, For** | **$H_2O$, $CO_2$, Glcn, For** | **$H_2O$, $CO_2$, Glcn, For** |
| | Ec | **$H_2O$, $CO_2$, Ace, $H^+$** | **$H_2O$, $CO_2$, Ace, $H^+$** | **$H_2O$, $CO_2$, $H^+$** |
| 2 | Ab | **$H_2O$, $CO_2$, Glcn, For** | **$H_2O$, $CO_2$, Glcn, For** | **$H_2O$, $CO_2$, Glcn, For** |
| | Ec | **$H_2O$, $CO_2$, Ace, $H^+$** | **$H_2O$, $CO_2$, Ace, $H^+$** | **$H_2O$, $CO_2$, $H^+$** |

The analyses were performed using the Python (`www.python.org`) programming language and the `cobrapy` library [5]. The libraries `numpy` [6] and `pandas` [7] were used both in preparation and analysis. The utilized genome-scale models of *A. baylyi* ADP1 and *E. coli* were `iAbaylyiv4` [8] and `iAF1260` [9], respectively. Growth associated maintenance (GAM) was set to 59.81 $mmol_{ATP}$ $g_{DW}^{-1}$ $h^{-1}$ (default of `iAF1260`) and non-growth associated maintenance (NGAM) to zero in both models. Setting NGAM to zero was necessary to keep the optimizations feasible when the knock-out strains were incapable of growing. The computational medium allowed for 8 mmol $g_{DW}^{-1}$ $h^{-1}$ glucose, oxygen as indicated above, and unlimited $Ca^{2+}$, $Cl^-$, $Co^{2-}$, $Fe^{2+}$, $K^+$, $Mg^{2+}$, $Mn^{2+}$, $MoO_4{}^{2-}$, $NH_4{}^+$, $HPO_4{}^-$, $Na^+$, $SO_4{}^{2-}$, $Zn^{2+}$, $H^+$, and $H_2O$, mimicking the medium used in both small- and larger-scale experiments (Table S6).

To correctly model for the effect of *ptsI* knock-out in *E. coli*, the reactions `GLCt2pp` (gluose-proton symport), `GLCabcpp` (glucose-ABC-transport system), and `GLCDpp` (glucose dehydrogenase) were permanently knocked out in iAF1260. Gluconate transport reaction from extracellular environment directly to cytoplasm (`TRANS-RXN-GLUCONATE`) was disabled in the `iAbaylyiv4` model and replaced by transport from extracellular space to periplasm in order to correctly model for *gntT* knock-out in *A. baylyi* ADP1. If these reactions were not removed, Ec$\Delta$*ptsI* (Ek) could have imported glucose through its other transport systems, and Ab$\Delta$*gntT* (Ak) could have imported gluconate through the passive transport of gluconate from extracellular environment to cytosol, rendering the knock-outs effectless.

The FBAs with an iterative scheme [1] were conducted with an additional step prepended to each iteration. Glucose importing exchange reactions (`EXF-GLC(E)` in `iAbaylyiv4` and `EX_glc_e_` in `iAF1260`) were maximized for first, prior to the actual growth rate maximization and subsequent gene-associated flux minimization. This was necessary to enable conversion of glucose to gluconate by Ab$\Delta$*gntT*. As Ab$\Delta$*gntT* was unable to grow on its own on glucose as the sole carbon source, the optimization procedure would not have resulted in glucose oxidation either. The maximized glucose import was then set as a fixed bound for the growth rate maximization and gene-associated flux minimization. As both knock-out strains Ab$\Delta$*gntT* and Ec$\Delta$*ptsI* were incapable of growing initially, it was necessary to incorporate the secreted metabolites into the computational medium regardless of the growth statuses, contrary to the original iterative FBA method [1]. The linking between `iAbaylyiv4` and `iAF160` extracellular metabolites was done manually.

### S2 Unstructured Kinetic Models

In this section the unstructured kinetic models referred to in the main text are derived. Considering a two-strain culture of total biomass concentration $X_\mathrm{T} = X_1 + X_2$ (mass volume$^{-1}$), the strain $i$'s proportion of total biomass, $X_i/X_\mathrm{T}$ (mass mass$^{-1}$), differentiated with respect to time $t$ is

$$\frac{\mathrm{d}\,(X_i/X_\mathrm{T})}{\mathrm{d}t} = \frac{1}{X_\mathrm{T}}\frac{\mathrm{d}X_i}{\mathrm{d}t} - \frac{X_i}{X_\mathrm{T}^2}\frac{\mathrm{d}X_\mathrm{T}}{\mathrm{d}t} \tag{S1}$$

Assuming the whole culture is growing or decaying ($\mathrm{d}X_\mathrm{T}/\mathrm{d}t \neq 0$) guarantees that eq S1 can be rewritten as

$$\frac{\mathrm{d}\,(X_i/X_\mathrm{T})}{\mathrm{d}t} = \frac{1}{X_\mathrm{T}}\frac{\mathrm{d}X_\mathrm{T}}{\mathrm{d}t}\left(\frac{\mathrm{d}X_i/\mathrm{d}t}{\mathrm{d}X_\mathrm{T}/\mathrm{d}t} - \frac{X_i}{X_\mathrm{T}}\right) \tag{S2}$$

The term $(\mathrm{d}X_i/\mathrm{d}t)\,(\mathrm{d}X_\mathrm{T}/\mathrm{d}t)^{-1}$ can now be interpreted as the equilibrium composition the consortium is adapting to. The task is then to identify this equilibrium state for the consortium at hand. Deriving an expression for $(\mathrm{d}X_i/\mathrm{d}t)\,(\mathrm{d}X_\mathrm{T}/\mathrm{d}t)^{-1}$ allows deducing whether the relative population abundances in the consortium adapt towards a fixed or a moving target.

Models for a consortium's strains need to derived first (Section S2.1) in order to identify the consortium's equilibrium composition (Section S2.2). The involved ordinary differential equations (ODE) systems are then solved for to investigate whether any assumptions involved significantly influenced the results (Section S2.3).

#### S2.1 Models of *A. baylyi* ADP1 and *E. coli*

For both *A. baylyi* ADP1 and *E. coli*, the specific growth rate $\mu_i$ (time$^{-1}$) on a substrate $i$ (mass volume$^{-1}$) was modeled as

$$\mu_i = \mu_i^* \frac{i}{K_i + i} \tag{S3}$$

where $\mu_i^*$ is the maximal specific growth rate (time$^{-1}$) and $K_i$ is an affinity constant (mass volume$^{-1}$). To make the subsequent equations more readable, the shorthand notation $i^* = i/(K_i + i)$ is used, which results in

$$\mu_i = \mu_i^* i^* \tag{S4}$$

The growth rates of *A. baylyi* ADP1 (a) and *E. coli* (e) are modeled as sums of growth rates on different substrates. *A. baylyi* ADP1 can oxidize glucose (G) to gluconate (N) and utilize acetate (A) and gluconate, whereas *E. coli* utilizes glucose and gluconate. Additionally, *E. coli* is assumed to produce acetate linearly upon utilization of glucose and gluconate. *E. coli* could utilize acetate simultaneously with glucose or gluconate, but net accumulation of acetate has been found to occur only at elevated acetate concentrations of approximately 10 mm or more [4]. Consequently, *E. coli* is considered only as a net producer of acetate in this analysis. Furthermore, a dilution factor $D$ (time$^{-1}$) and glucose feed $f_G$ (mass volume$^{-1}$ time$^{-1}$) are considered in order to not limit the analysis only to pure batch cultivations. In batch cultures both $D = 0$ and $f_G = 0$ apply, whereas in fed-batches and continuous operations $D > 0$ and $f_G > 0$ are found instead.

Denoting then *A. baylyi* ADP1 concentration (mass volume$^{-1}$) as $X_a$, and *E. coli* concentration (mass volume$^{-1}$) as $X_e$, the biomass concentration time derivatives are

$$\frac{dX_a}{dt} = \left(\mu_{Aa}^* A^* + \mu_{Na}^* N^* - D\right) X_a \tag{S5}$$

$$\frac{dX_e}{dt} = \left(\mu_{Ge}^* G^* + \mu_{Ne}^* N^* - D\right) X_e \tag{S6}$$

In order to then identify the equilibrium proportion of *A. baylyi* ADP1 in an *A. baylyi* ADP1:*E. coli* consortium, $(dX_a/dt)\,(dX_T/dt)^{-1}$, and to detect whether it is a constant or a variable, the terms $G^*$, $N^*$, and $A^*$ are required. To solve for these, the reactions catalyzed by *A. baylyi* and *E. coli* along with their rates have to be defined first. *A. baylyi* ADP1 can catalyze the following reactions:

$$\mathrm{G} \xrightarrow{\mathrm{R_1}} \mathrm{Y_{GN}\,N}$$

$$\mathrm{N} \xrightarrow{\mathrm{R_3}} \mathrm{Y_{Na}\,X_a}$$

$$\mathrm{A} \xrightarrow{\mathrm{R_5}} \mathrm{Y_{Aa}\,X_a}$$

which correspond to glucose oxidation to gluconate ($R_1$), growth on gluconate ($R_3$), and growth on acetate ($R_5$). The yield coefficients $Y_{sp}$ denote the mass of $p$ produced per a mass of $s$ consumed. *E. coli*, on the other hand, can catalyze the reactions

$$\mathrm{G} \xrightarrow{\mathrm{R_2}} \mathrm{Y_{Ge}\,X_e + Y_{GA}\,A}$$

$$\mathrm{N} \xrightarrow{\mathrm{R_4}} \mathrm{Y_{Ne}\,X_e + Y_{NA}\,A}$$

which stand for growth on glucose ($R_2$) and gluconate ($R_4$) along with linear acetate excretion. The reaction rates $R$ ($\mathrm{mass_s}$ $\mathrm{volume}^{-1}$ $\mathrm{time}^{-1}$) for these five reactions are then written based on the reaction substrates $s \in (G, N, A)$:

$$R_1 = q^*_{\mathrm{Ga}} X_{\mathrm{a}} G^* \tag{S7}$$

$$R_2 = \mu^*_{\mathrm{Ge}} X_{\mathrm{e}} Y^{-1}_{\mathrm{Ge}} G^* \tag{S8}$$

$$R_3 = \mu^*_{\mathrm{Na}} X_{\mathrm{a}} Y^{-1}_{\mathrm{Na}} N^* \tag{S9}$$

$$R_4 = \mu^*_{\mathrm{Ne}} X_{\mathrm{e}} Y^{-1}_{\mathrm{Ne}} N^* \tag{S10}$$

$$R_5 = \mu^*_{\mathrm{Aa}} X_{\mathrm{a}} Y^{-1}_{\mathrm{Aa}} A^* \tag{S11}$$

where $\mu^*_{ij}$ is the maximal specific growth rate of strain $j$ on substrate $i$ ($\mathrm{time}^{-1}$), and $q^*_{ij}$ is the maximal specific reaction rate ($\mathrm{mass_i}$ $\mathrm{mass_j}^{-1}$ $\mathrm{time}^{-1}$) of $i$ catalyzed by $j$. Figure S1 illustrates the five-reaction unstructured kinetic model of an *A. baylyi* ADP1:*E. coli* consortium, and Table S2 shows how the five reactions are used in modeling the four consortia.

Considering glucose feed $f_{\mathrm{G}}$, dilution factor $D$, and the defined reactions rates, mass balances for glucose, gluconate, and acetate are written as

$$\frac{\mathrm{d}G}{\mathrm{d}t} = f_{\mathrm{G}} - R_1 - R_2 - DG \tag{S12}$$

$$\frac{\mathrm{d}N}{\mathrm{d}t} = Y_{\mathrm{GN}} R_1 - R_3 - R_4 - DN \tag{S13}$$

$$\frac{\mathrm{d}A}{\mathrm{d}t} = Y_{\mathrm{GA}} R_2 + Y_{\mathrm{NA}} R_4 - R_5 - DA \tag{S14}$$

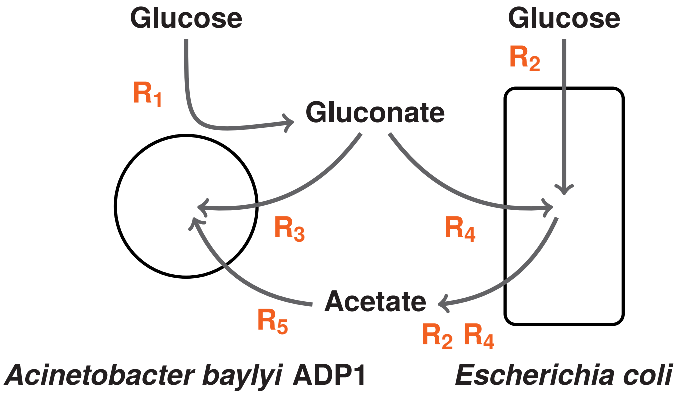

Figure S1: Reactions in an unstructured kinetic model of an *A. baylyi* ADP1:*E. coli* consortium. Table S2 shows how the reactions are used in modeling the four consortia.

In batch cultures both the glucose feed $f_G$ and the dilution factor $D$ equal zero.

Given the reaction rate eqs S7, S8, S9, S10, and S11, and the mass balance eqs S12, S13, and S14, the terms $G^*$, $N^*$, and $A^*$ are solved to be

$$G^* = \frac{f_G + (\mathrm{d}G/\mathrm{d}t) - DG}{q^*_{Ga}X_a + \mu^*_{Ge}X_e Y^{-1}_{Ge}} \tag{S15}$$

$$N^* = \frac{Y_{GN} q^*_{Ga} X_a G^* + (\mathrm{d}N/\mathrm{d}t) - DN}{\mu^*_{Na}X_a Y^{-1}_{Na} + \mu^*_{Ne}X_e Y^{-1}_{Ne}} \tag{S16}$$

$$A^* = \frac{Y_{GA}\mu^*_{Ge}X_e Y^{-1}_{Ge} G^* + Y_{NA}\mu^*_{Ne}X_e Y^{-1}_{Ne} N^* + (\mathrm{d}A/\mathrm{d}t) - DA}{\mu^*_{Aa}X_a Y^{-1}_{Aa}} \tag{S17}$$

#### S2.2 Optimal consortium compositions

With the terms $G^*$, $N^*$, and $A^*$ (eqs S15, S16, and S17) at hand, the *A. baylyi* ADP1 and *E. coli* biomass time derivatives (eqs S5 and S6) can be defined to identify the optimal *A. baylyi* ADP1 proportion, $(\mathrm{d}X_a/\mathrm{d}t)\,(\mathrm{d}X_T/\mathrm{d}t)^{-1}$, in four consortia involving *A. baylyi* ADP1 and *E. coli*, namely AbΔ*gntT*:EcΔ*ptsI*, AbΔ*gntT*:Ec, Ab:EcΔ*ptsI*, and Ab:Ec. To facilitate derivation, the following shorthand notation is used for glucose, gluconate, and acetate: $i' = (\mathrm{d}i/\mathrm{d}t) - Di$.

**AbΔ*gntT*:EcΔ*ptsI*** In AbΔ*gntT*:EcΔ*ptsI*, the reactions $R_1$, $R_4$, and $R_5$ are active, corresponding to glucose oxidation by *A. baylyi* ADP1 to gluconate, growth on gluconate and simultaneous excretion of acetate by *E. coli*, and growth on acetate by *A. baylyi*

Table S2: Reactions used in modeling the four *A. baylyi* ADP1:*E. coli* consortia, Ab$\Delta$*gntT*:Ec$\Delta$*ptsI*, Ab$\Delta$*gntT*:Ec, Ab:Ec$\Delta$*ptsI*, and Ab:Ec. A reaction's presence in a particular consortium is marked with + or 0, and - is used to designate a reaction's absence. 0 is used to show which reactions have been omitted in Section S2.2 in order to simplify equations. However, both + and 0 reactions were used in solving the initial value problems (Section S2.3).

| Reaction | Ab$\Delta$*gntT*:Ec$\Delta$*ptsI* | Ab$\Delta$*gntT*:Ec | Ab:Ec$\Delta$*ptsI* | Ab:Ec |
|---|---|---|---|---|
| 1 | + | 0 | + | + |
| 2 | - | + | - | + |
| 3 | - | - | + | + |
| 4 | + | 0 | + | 0 |
| 5 | + | + | 0 | 0 |

ADP1. Given these reactions and the equations defined above, the equilibrium proportion of Ab$\Delta$*gntT* becomes

$$\frac{\mathrm{d}X_\mathrm{a}/\mathrm{d}t}{\mathrm{d}X_\mathrm{T}/\mathrm{d}t} = \frac{\mu^*_\mathrm{Aa} X_\mathrm{a} A^* - D X_\mathrm{a}}{\mu^*_\mathrm{Aa} X_\mathrm{a} A^* - D X_\mathrm{a} + \mu^*_\mathrm{Ne} X_\mathrm{e} N^* - D X_\mathrm{e}} \tag{S18}$$

which expands to

$$\frac{\mathrm{d}X_\mathrm{a}/\mathrm{d}t}{\mathrm{d}X_\mathrm{T}/\mathrm{d}t} = \frac{Y_\mathrm{Aa} Y_\mathrm{NA} \left(Y_\mathrm{GN} \left(f_\mathrm{G} + G'\right) + N'\right) + Y_\mathrm{Aa} A' - D X_\mathrm{a}}{\left(Y_\mathrm{Aa} Y_\mathrm{NA} + Y_\mathrm{Ne}\right) \left(Y_\mathrm{GN} \left(f_\mathrm{G} + G'\right) + N'\right) + Y_\mathrm{Aa} A' - D X_\mathrm{T}} \tag{S19}$$

Assuming only that acetate reaches a steady state of constant concentration, $\mathrm{d}A/\mathrm{d}t \approx 0$, and that the dilution rate is neglicible (equals zero in batch, only small in fed-batch), $D \approx 0$, eq S19 simplifies to

$$\frac{\mathrm{d}X_\mathrm{a}/\mathrm{d}t}{\mathrm{d}X_\mathrm{T}/\mathrm{d}t} = \frac{Y_\mathrm{Aa} Y_\mathrm{NA}}{Y_\mathrm{Aa} Y_\mathrm{NA} + Y_\mathrm{Ne}} \tag{S20}$$

Further assuming that the yield coefficients are constants, eq S20 states that there is a uniquely defined, constant equilibrium composition the consortium adapts to. Therefore, the double knock-out Ab$\Delta$*gntT*:Ec$\Delta$*ptsI* consortium with the interconnected carbon cross-feeding is expected to be stable.

**Ab$\Delta$*gntT*:Ec** As for Ab$\Delta$*gntT*:Ec, *A. baylyi* ADP1 oxidizes glucose to gluconate and *E. coli* grows on glucose and gluconate and excretes acetate while doing so. *A. baylyi* ADP1 grows on the acetate secreted by *E. coli*, but as it cannot import gluconate, the reaction

$R_3$ is set to zero. In deriving the optimal strain proportion expression, reactions $R_1$ and $R_4$ (glucose oxidation by *A. baylyi* ADP1, gluconate utilization by *E. coli*) are also set to zero in order to simplify the analysis. Given the simplifications, AbΔ*gntT*:Ec becomes a commensialistic consortium, in which AbΔ*gntT* benefits from Ec without having an influence on Ec. The equilibrium proportion of *A. baylyi* ADP1 in AbΔ*gntT*:Ec is then

$$\frac{\mathrm{d}X_\mathrm{a}/\mathrm{d}t}{\mathrm{d}X_\mathrm{T}/\mathrm{d}t} = \frac{\mu^*_\mathrm{Aa} X_\mathrm{a} A^* - DX_\mathrm{a}}{\mu^*_\mathrm{Aa} X_\mathrm{a} A^* - DX_\mathrm{a} + \mu^*_\mathrm{Ge} X_\mathrm{e} G^* - DX_\mathrm{e}} \tag{S21}$$

which expands to

$$\frac{\mathrm{d}X_\mathrm{a}/\mathrm{d}t}{\mathrm{d}X_\mathrm{T}/\mathrm{d}t} = \frac{Y_\mathrm{Aa} Y_\mathrm{GA} \left(f_\mathrm{G} + G'\right) + Y_\mathrm{Aa} A' - DX_\mathrm{a}}{\left(Y_\mathrm{Aa} Y_\mathrm{GA} + Y_\mathrm{Ge}\right)\left(f_\mathrm{G} + G'\right) + Y_\mathrm{Aa} A' - DX_\mathrm{T}} \tag{S22}$$

Assuming again that both $\mathrm{d}A/\mathrm{d}t \approx 0$ and $D \approx 0$ apply, eq S22 simplifies to

$$\frac{\mathrm{d}X_\mathrm{a}/\mathrm{d}t}{\mathrm{d}X_\mathrm{T}/\mathrm{d}t} = \frac{Y_\mathrm{Aa} Y_\mathrm{GA}}{Y_\mathrm{Aa} Y_\mathrm{GA} + Y_\mathrm{Ge}} \tag{S23}$$

Like with AbΔ*gntT*:EcΔ*ptsI*, the equilibrium proportion simplified to a function of yield coefficients only, suggesting stability. However, numerical integrations (Section S2.3) of the ODE system indicated that the assumption of constant acetate concentration, $\mathrm{d}A/\mathrm{d}t \approx 0$, does not hold as well as in AbΔ*gntT*:EcΔ*ptsI*. Considering that growth and acetate production of *E. coli* is entirely independent of *A. baylyi* ADP1, it is logical to expect that *E. coli* is capable of accumulating acetate beyond *A. baylyi* ADP1's needs, collapsing the simple eq S23 back to the more complex eq S22 as $\mathrm{d}A/\mathrm{d}t \approx 0$ does not apply. Additionally, the expressions and consequently dynamics would be more complex if glucose oxidation to gluconate by *A. baylyi* ADP1 ($R_1$) and gluconate utilization by *E. coli* ($R_4$) were considered.

**Ab:EcΔ*ptsI*** The Ab:EcΔ*ptsI* consortium was modeled with the reactions $R_1$, $R_3$, and $R_4$, which corresponded to glucose oxidation to gluconate by *A. baylyi* ADP1 and growth on gluconate by both *A. baylyi* ADP1 and *E. coli*. However, to simplify derivation of the optimal strain proportions, the yield from gluconate to acetate by *E. coli* is set to equal zero. Likewise, acetate utilization by *A. baylyi* ADP1 was set to zero as none was produced by *E. coli*. With these simplifications, Ab:EcΔ*ptsI* has cooperator-cheater dynamics, in which EcΔ*ptsI* benefits from and also has a negative effect on Ab as they both grow on gluconate. Taking these into account, the equilibrium proportion of *A. baylyi* ADP1 in Ab:EcΔ*ptsI* can be written as

$$\frac{\mathrm{d}X_\mathrm{a}/\mathrm{d}t}{\mathrm{d}X_\mathrm{T}/\mathrm{d}t} = \frac{\mu^*_\mathrm{Na} X_\mathrm{a} N^* - DX_\mathrm{a}}{\mu^*_\mathrm{Na} X_\mathrm{a} N^* - DX_\mathrm{a} + \mu^*_\mathrm{Ne} X_\mathrm{e} N^* - DX_\mathrm{e}} \tag{S24}$$

Assuming negligible dilution, $D \approx 0$, leads in turn to

$$\frac{\mathrm{d}X_\mathrm{a}/\mathrm{d}t}{\mathrm{d}X_\mathrm{T}/\mathrm{d}t} = \frac{\mu^*_\mathrm{Na}X_\mathrm{a}}{\mu^*_\mathrm{Na}X_\mathrm{a} + \mu^*_\mathrm{Ne}X_\mathrm{e}} \tag{S25}$$

Now in contrast to both AbΔ*gntT*:EcΔ*ptsI* and AbΔ*gntT*:Ec, the optimal *A. baylyi* ADP1 proportion is found to be a variable dependent on the biomass concentrations themselves. Again, the expression would be more complex if acetate excretion by *E. coli* and utilization by *A. baylyi* ADP1 were taken into account.

**Ab:Ec** The wild-type Ab:Ec consortium includes all reactions from $\mathrm{R}_1$ to $\mathrm{R}_5$. However, to simplify derivation of the optimal strain proportions, the reactions $\mathrm{R}_4$ and $\mathrm{R}_5$ corresponding to gluconate utilization by *E. coli* and acetate utilization by *A. baylyi* ADP1 are omitted. The consortium involves direct competition between Ab and Ec, as they both utilize glucose. The equilibrium proportion of *A. baylyi* ADP1 is then

$$\frac{\mathrm{d}X_\mathrm{a}/\mathrm{d}t}{\mathrm{d}X_\mathrm{T}/\mathrm{d}t} = \frac{\mu^*_\mathrm{Na}X_\mathrm{a}N^* - DX_\mathrm{a}}{\mu^*_\mathrm{Na}X_\mathrm{a}N^* - DX_\mathrm{a} + \mu^*_\mathrm{Ge}X_\mathrm{e}G^* - DX_\mathrm{e}} \tag{S26}$$

or equivalently

$$\frac{\mathrm{d}X_\mathrm{a}/\mathrm{d}t}{\mathrm{d}X_\mathrm{T}/\mathrm{d}t} = \frac{Y_\mathrm{Na}Y_\mathrm{GN}q^*_\mathrm{Ga}X_\mathrm{a}\frac{f_\mathrm{G}+G'}{q^*_\mathrm{Ga}X_\mathrm{a}+\mu^*_\mathrm{Ge}X_\mathrm{e}Y^{-1}_\mathrm{Ge}} + Y_\mathrm{Na}N' - DX_\mathrm{a}}{\left(Y_\mathrm{Na}Y_\mathrm{GN}q^*_\mathrm{Ga}X_\mathrm{a} + \mu^*_\mathrm{Ge}X_\mathrm{e}\right)\frac{f_\mathrm{G}+G'}{q^*_\mathrm{Ga}X_\mathrm{a}+\mu^*_\mathrm{Ge}X_\mathrm{e}Y^{-1}_\mathrm{Ge}} + Y_\mathrm{Na}N' - DX_\mathrm{T}} \tag{S27}$$

Now assuming negligible dilution, $D \approx 0$, and constant gluconate concentration, $\mathrm{d}N/\mathrm{d}t \approx 0$, results in

$$\frac{\mathrm{d}X_\mathrm{a}/\mathrm{d}t}{\mathrm{d}X_\mathrm{T}/\mathrm{d}t} = \frac{Y_\mathrm{Na}Y_\mathrm{GN}q^*_\mathrm{Ga}X_\mathrm{a}}{Y_\mathrm{Na}Y_\mathrm{GN}q^*_\mathrm{Ga}X_\mathrm{a} + \mu^*_\mathrm{Ge}X_\mathrm{e}} \tag{S28}$$

Like in Ab:EcΔ*ptsI*, the optimal strain proportions are found to be a variable dependent on the biomass concentrations themselves. As with AbΔ*gntT*:Ec and Ab:EcΔ*ptsI* alike, the expressions would be more complex if all the relevant reactions were fully taken into account.

**Conclusion** Based on these simple kinetic analyses, it is concluded that the interconnected carbon cross-feeding present in AbΔ*gntT*:EcΔ*ptsI* best guarantees that a consortium's composition moves towards a unique and stable target. The other consortia corresponding to commensialism, cooperator-cheater, and competition dynamics had equilibrium proportions that were susceptible to either prevailing biomass concentrations or acetate fluctuations.

Table S3: Initial value problem parameters. $X_a$ and $X_e$ stand for initial biomass concentrations of *A. baylyi* ADP1 and *E. coli*, respectively. $G$ is the initial glucose concentration. $\beta$ is the exponential coefficient of the feed flow-rate $F$ in fed-batches. Zero exponential $\beta = 0$ corresponds to constant feed rate.

| Batch | $X_a$ / g L$^{-1}$ | $X_e$ / g L$^{-1}$ | $G$ / g L$^{-1}$ |
|---|---|---|---|
| 1 | 0.050 | 0.050 | 9.00 |
| 2 | 0.050 | 0.050 | 18.00 |
| 3 | 0.050 | 0.050 | 36.00 |
| 4 | 0.033 | 0.067 | 18.00 |
| 5 | 0.017 | 0.083 | 18.00 |
| 6 | 0.010 | 0.090 | 18.00 |
| **Fed-batch** | $X_a$ / g L$^{-1}$ | $X_e$ / g L$^{-1}$ | $\beta$ / h$^{-1}$ |
| 1 | 0.050 | 0.050 | 0.00 |
| 2 | 0.050 | 0.050 | 0.10 |
| 3 | 0.050 | 0.050 | 0.20 |
| 4 | 0.033 | 0.067 | 0.10 |
| 5 | 0.017 | 0.083 | 0.10 |
| 6 | 0.010 | 0.090 | 0.10 |

#### S2.3 Initial value problem solutions

The behaviour of the derived *A. baylyi* ADP1:*E. coli* consortium models was inspected numerically in both batch and fed-batch modes. The batch simulations had the same initial glucose concentrations and inoculations ratios as the conducted small-scale experiments. All simulations were performed in Python using the `scipy` library's [10] `solve_ivp` function with the default `RK45` method. The libraries `numpy` [6] and `pandas` [7] were also used in preparation and analysis.

Taking the five reactions as shown in Table S2 (both + and 0), and the relevant mass balances in eqs S5, S6, S12, S13, and S14, the four ODE systems corresponding to the four *A. baylyi* ADP1:*E. coli* consortia were solved in both batch and fed-batch mode with varying initial substrate concentrations, feed parameters, and inoculation ratios as described in Table S3. The fed-batches of volume $V = 1$ L had always an initial glucose concentration of 0.1 g L$^{-1}$ and a feed with

1. Glucose concentration $S_0 = 520\,\mathrm{g\,L^{-1}}$
2. Initial flow-rate $F_0 = 0.001\,\mathrm{L\,h^{-1}}$
3. Maximal flow-rate $F_{max} = 0.01\,\mathrm{L\,h^{-1}}$
4. Time $t$ dependency of flow-rate $F = \min\left(F_0 e^{\beta t}, F_{max}\right)$ characterized by an exponential coefficient $\beta$ (time$^{-1}$), which was varied between fed-batches.

Table S4: Yield coefficients $Y_{sp}$ (mass of $p$ produced per mass of $s$ consumed) and maximal specific growth rates $\mu^*_{sx}$ (rate of $x$ growing on $s$) used in solving the initial value problems. *A. baylyi* ADP1 biomass yield on gluconate ($Y_{\mathrm{Na}}$) was estimated by considering that an engineered *A. baylyi* ADP1 expressing PykF (pyruvate kinase from *E. coli*), which was found to have similar biomass yield as the wild-type [11], accumulated 0.30 g L$^{-1}$ biomass given 10 mᴍ glucose in a minimal medium [12]. The biomass yield on glucose was then translated to gluconate by considering the ratio of their molar masses. Due to lack of more precise information, *E. coli* biomass yield on gluconate ($Y_{\mathrm{Ne}}$) was assumed to be similar to yield on glucose ($Y_{\mathrm{Ge}}$) [13], but divided by the ratio of gluconate's molar mass to glucose's molar mass. The same was applied to *E. coli*'s maximal specific growth rate on gluconate ($\mu^*_{\mathrm{Ne}}$).

| Yields $Y_{sp}$ | $\mathrm{g_p\,g_s^{-1}}$ | Source |
|---|---|---|
| $Y_{\mathrm{Na}}$ | 0.32 | [11, 12] |
| $Y_{\mathrm{Aa}}$ | 0.50 | [12] |
| $Y_{\mathrm{Ge}}$ | 0.50 | [13] |
| $Y_{\mathrm{Ne}}$ | 0.46 | [13], molar masses |
| $Y_{\mathrm{GN}}$ | 1.09 | Molar masses |
| $Y_{\mathrm{GA}}$ | 0.10 | [4] |
| $Y_{\mathrm{NA}}$ | 0.15 | [4] |
| **Maximal specific growth rates $\mu^*_{sx}$** | **$\mathrm{h^{-1}}$** | **Source** |
| $\mu^*_{\mathrm{Aa}}$ | 0.69 | [12] |
| $\mu^*_{\mathrm{Na}}$ | 0.18 | [11] |
| $\mu^*_{\mathrm{Ge}}$ | 0.66 | [13] |
| $\mu^*_{\mathrm{Ne}}$ | 0.61 | [13], molar masses |

In fed-batches the dilution factor $D$ was then $D = F/V$ and glucose feed $f_{\mathrm{G}} = S_0F/V$. The yield coefficients $Y_{sp}$ ($\mathrm{mass_p\,mass_s^{-1}}$) and maximal specific rates $\mu^*_{ij}$ ($\mathrm{time^{-1}}$) used in the kinetic models are shown in Table S4. All affinity constants $K_i$ were taken to be 0.05 g L$^{-1}$ as in [13], and *A. baylyi* ADP1's maximal specific glucose oxidation rate $q^*_{\mathrm{Ga}}$ was assumed to be 2.0 $\mathrm{g_G\,g_a^{-1}\,h^{-1}}$.

The obtained solutions to the initial value problems are summarized in Figures S2 and S3 for batch and fed-batch modes, respectively. It was shown in Section S2.2, that both Ab$\Delta$*gntT*:Ec$\Delta$*ptsI* and Ab$\Delta$*gntT*:Ec have unique and constant optimum strain proportions given that acetate concentration $A$ is constant and dilution $D$ is negligible. Figures S2 and S3 demonstrate that the equilibrium proportion of *A. baylyi* ADP1 $(\mathrm{d}X_\mathrm{a}/\mathrm{d}t)\,(\mathrm{d}X_\mathrm{T}/\mathrm{d}t)^{-1}$ in Ab$\Delta$*gntT*:Ec$\Delta$*ptsI* (eq S20) remained stable in both batches and fed-batches with only small deviations caused by the barely visible initial acetate build-ups. In Ab$\Delta$*gntT*:Ec,

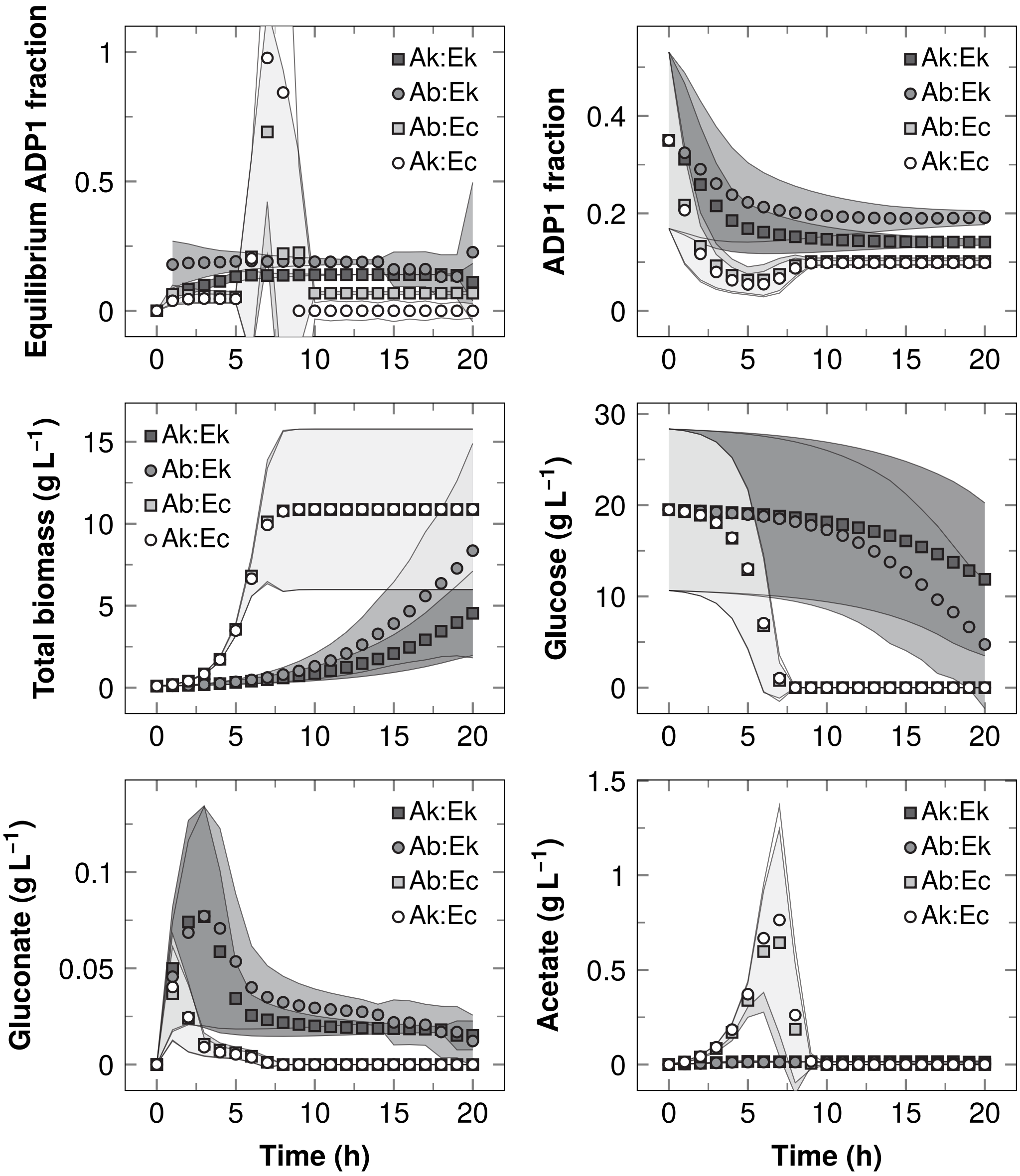

Figure S2: Initial value problem solutions in batch mode. Ab stands for *A. baylyi* ADP1, Ak for Ab$\Delta$*gntT*, Ec for *E. coli*, and Ek for Ec$\Delta$*ptsI*. The reported data are means of the six simulations with different initial conditions (Table S3). The variability between simulations is shown as error bands corresponding to sample standard deviations. The legend in glucose panel was omitted for clarity.

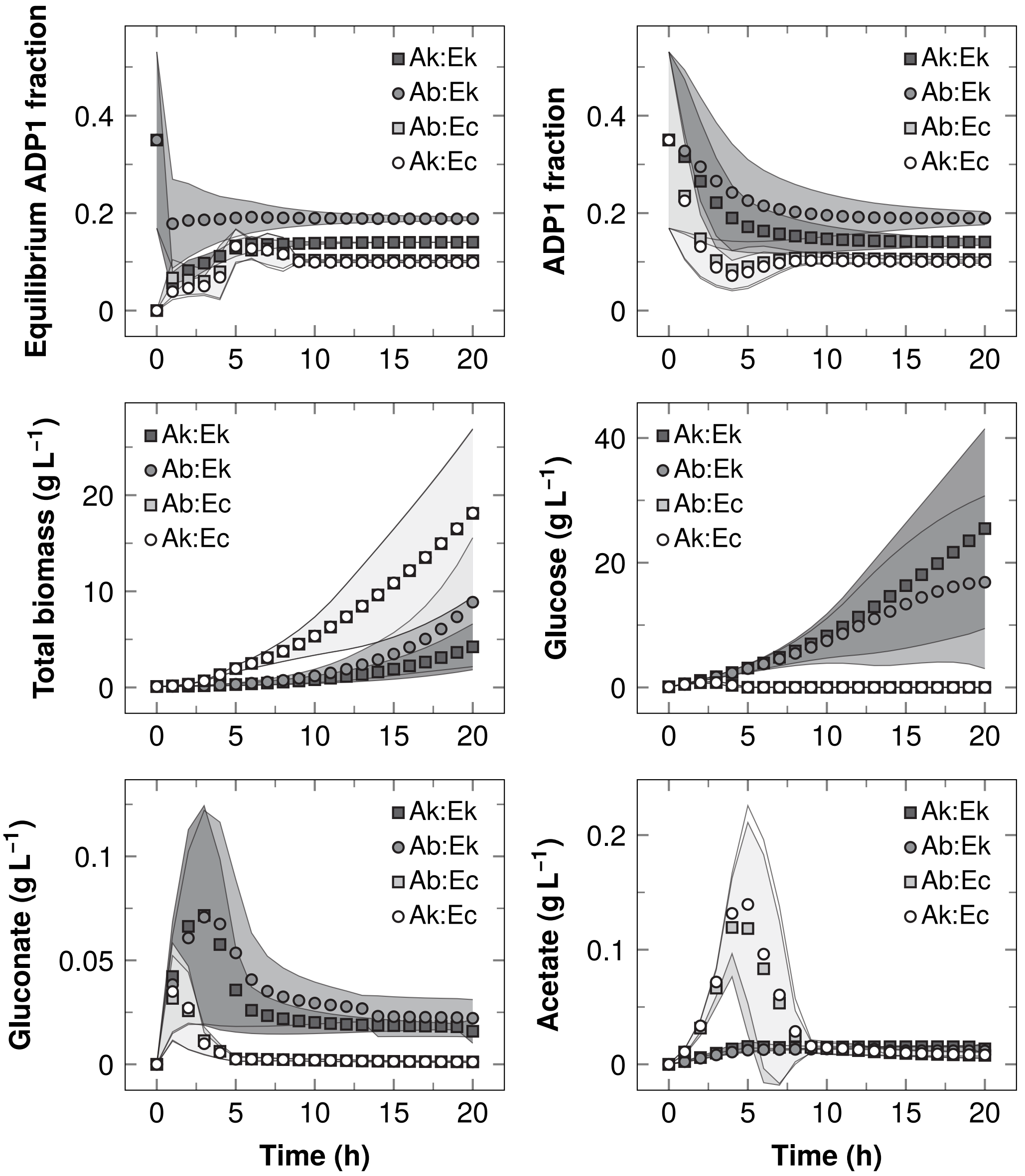

Figure S3: Initial value problem solutions in fed-batch mode. Ab stands for *A. baylyi* ADP1, Ak for Ab$\Delta$*gntT*, Ec for *E. coli*, and Ek for Ec$\Delta$*ptsI*. The reported data are means of the six simulations with different initial conditions (Table S3). The variability between simulations is shown as error bands corresponding to sample standard deviations.

however, acetate accumulated ($\mathrm{d}A/\mathrm{d}t \neq 0$) markedly both in batches and fed-batches as growth and acetate secretion by *E. coli* was not in any way limited by *A. baylyi* ADP1. As a consequence, the equilibrium proportion of *A. baylyi* ADP1 did not remain constant, but displayed an oscillation. The actual *A. baylyi* ADP1 fraction of total biomass ($X_\mathrm{a}/X_\mathrm{T}$) also varied less in AbΔ*gntT*:EcΔ*ptsI* than in AbΔ*gntT*:Ec.

As for Ab:EcΔ*ptsI* and Ab:Ec, it was shown that the equilibrium strain proportions were a function of the biomass concentrations themselves (eqs S25 and S28), and consequently neither performed as steadily as AbΔ*gntT*:EcΔ*ptsI*; Ab:Ec was in general similar to AbΔ*gntT*:Ec with an oscillation in both equilibrium and actual *A. baylyi* ADP1 fractions, and Ab:EcΔ*ptsI* displayed more variance in *A. baylyi* ADP1 fractions than AbΔ*gntT*:EcΔ*ptsI*.

### S3 mScarlet Gene Cassette Construction

This section describes how the mScarlet expression cassette was constructed using the insert ordered from GenScript, USA, and a gene cassette [14] overwriting the neutral ACIAD3381-locus (*poxB*, pyruvate dehydrogenase). The mScarlet insert consisted of

1. MunI (MfeI) recognition site
2. BBa_J23100 promoter present also in the sfGFP cassette [15]
3. Synthetic ribosome binding site (RBS) present also in the sfGFP cassette [15], but shortened 3 bp from 3' end to accommodate the NdeI recognition site
4. NdeI recognition site which supplied the start codon
5. 6His-tag [16]
6. Gly-Ser-Gly-linker between 6His-tag and mScarlet
7. mScarlet sequence based on the amino acid sequence [17] codon optimized for *E. coli* by GenScript, start codon excluded
8. Two stop codons
9. XhoI recognition site.

The insert is illustrated in Figure S4 and its sequences are shown in Table S5. The pUC57 plasmid provided by GenScript with the mScarlet insert was first amplified by growing its initial *E. coli* host in lysogeny broth (LB). Using MunI and XhoI restriction enzymes (Fermentas, Lithuania), both the pUC57 mScarlet and the ADP1 gene cassette [14] were digested. The mScarlet insert was then ligated into the plasmid backbone using T4 DNA Ligase in T4 DNA Ligase Buffer (Thermo Scientific, USA).

The ligated plasmid product with the mScarlet insert was transformed by electroporation into *E. coli* XL1 (Stratagen, USA) for amplification and extraction. Cells were made electrocompetent, and the transformation was conducted using a MicroPulser electroporator (Bio-Rad, USA) with the Eco-1 program. The electroporated cell suspension was

Table S5: Sequences of the mScarlet construct inserted into a gene cassette [14] overwriting the neutral ACIAD0544 locus (*poxB*, pyruvate dehydrogenase). The shown mScarlet sequence is based on the amino acid sequence published in the original work [17]. The actual coding sequence was codon optimized for *E. coli* by GenScript.

| Part | Sequence (5′ → 3′) | bp | Source |
|---|---|---|---|
| **Upstream** | | | |
| MunI | CAATTG | 6 | |
| Promoter | TTGACGGCTAGCTCAGTCCTAGGTACAGTGCTAGC | 35 | [15] |
| RBS | TACTAGAGAAATCAAATTAAGGAGGTAAG | 29 | [15] |
| NdeI (1 to 3) | CAT | 3 | |
| **Coding sequence** | | | |
| NdeI (4 to 6) | ATG | 3 | |
| His-tag | CATCATCATCATCATCAC | 18 | |
| GSG-linker | GGTTCTGGT | 9 | |
| mScarlet | GTGAGCAAGGGCGAGGCAGTGATCAAGGAGTTCATGCGGTTC<br>AAGGTGCACATGGAGGGCTCCATGAACGGCCACGAGTTCGAG<br>ATCGAGGGCGAGGGCGAGGGCCGCCCCTACGAGGGCACCCAG<br>ACCGCCAAGCTGAAGGTGACCAAGGGTGGCCCCCTGCCCTTC<br>TCCTGGGACATCCTGTCCCCTCAGTTCATGTACGGCTCCAGG<br>GCCTTCACCAAGCACCCCGCCGACATCCCCGACTACTATAAG<br>CAGTCCTTCCCCGAGGGCTTCAAGTGGGAGCGCGTGATGAAC<br>TTCGAGGACGGCGGCGCCGTGACCGTGACCCAGGACACCTCC<br>CTGGAGGACGGCACCCTGATCTACAAGGTGAAGCTCCGCGGC<br>ACCAACTTCCCTCCTGACGGCCCCGTAATGCAGAAGAAGACA<br>ATGGGCTGGGAAGCGTCCACCGAGCGGTTGTACCCCGAGGAC<br>GGCGTGCTGAAGGGCGACATTAAGATGGCCCTGCGCCTGAAG<br>GACGGCGGCCGCTACCTGGCGGACTTCAAGACCACCTACAAG<br>GCCAAGAAGCCCGTGCAGATGCCCGGCGCCTACAACGTCGAC<br>CGCAAGTTGGACATCACCTCCCACAACGAGGACTACACCGTG<br>GTGGAACAGTACGAACGCTCCGAGGGCCGCCACTCCACCGGC<br>GGCATGGACGAGCTGTACAAG | 693 | [17] |
| Stop | TAA | 3 | |
| **Downstream** | | | |
| Stop | TAA | 3 | |
| XhoI | CTCGAG | 6 | |
| **Total** | | 808 | |

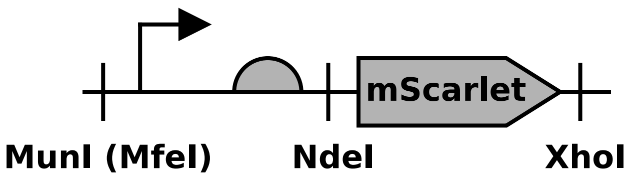

Figure S4: The mScarlet construct inserted into a gene cassette [14, see Figure 2] overwriting the neutral ACIAD0544 locus (*poxB*, pyruvate dehydrogenase). The promoter BBa_J23100 and ribosome binding site were from [15]. See Table S5 for the sequences. Figure was prepared with DNAplotlib [18].

plated on a LA plate (15 g L$^{-1}$ agar, 10 g L$^{-1}$ tryptone, 5 g L$^{-1}$ yeast extract, 1 g L$^{-1}$ NaCl, and 10 g L$^{-1}$ glucose) with 25 mg L$^{-1}$ chloramphenicol. The colonies of transformed *E. coli* XL1 that grew on the plate were bright red even to naked eye. A bright red colony was selected for amplification, and the mScarlet-carrying gene cassette plasmid was extracted using GeneJET Plasmid MiniPrep Kit (Thermo Scientific). Integration of the resulting mScarlet gene cassette into *A. baylyi* ADP1 has been described in the main text.

### S4 sfGFP Integrations

This section briefly describes how the sfGFP-carrying Burden Monitor expression cassette [15] was integrated into both *E. coli* and *E. coli* Δ*ptsI* using the conditional-replication, integration, and modular plasmids as described in the original publication [19]. Prior to all transformations cells were cultured in LB medium at 37 °C with 300 RPM shaking and made electrocompetent, and all transformations were conducted using a MicroPulser electroporator (Bio-Rad) with the Eco-1 program. *E. coli* was first transformed with the Int (integrase) expressing helper plasmid pAH123. The pAH123 helper plasmid carried also an ampicillin resistance gene. After transformation, *E. coli* was resistant to ampicillin as expected and grew on the selective LA plates containing 100 mg L$^{-1}$ ampicillin.

The pAH123 carrying *E. coli* strains were then further transformed with pBM (Burden Monitor phi80 version plasmid), yielding gentamicin-resistant strains. Immediately after electroporation the cells were incubated in LB medium at 37 °C for 1 h and furthermore at 42 °C for 30 min. The higher temperatures drove Int synthesis and inhibited helper plasmid replication [19]. The electroporated cell suspensions were then transferred to LA plates with 15 mg L$^{-1}$ gentamicin, and visibly green colonies were formed within few days of incubation at 37 °C. Curing of the helper plasmid pAH123 was confirmed by transferring colonies of the pBM-transformed strains to ampicillin containing LA plates. No growth was observed on the ampicillin plates.

Table S6: Composition of the defined medium used in experiments and precultivations. The medium is based on a mineral salt medium [20]. A carbon source was always added to medium prior to use as described in main text.

| Component | $mg\,L^{-1}$ | Component | $mg\,L^{-1}$ |
|---|---|---|---|
| $K_2HPO_4$ | 3880 | $ZnSO_4 \cdot 7\,H_2O$ | 2 |
| $NaH_2PO_4$ | 1630 | $CaCl_2 \cdot 2\,H_2O$ | 1 |
| $(NH_4)_2SO_4$ | 2000 | $MnCl_2 \cdot 2\,H_2O$ | 1 |
| $MgCl_2 \cdot 6\,H_2O$ | 100 | $CoCl_2 \cdot 6\,H_2O$ | 0.4 |
| EDTA | 10 | $CuSO_4 \cdot 5\,H_2O$ | 0.2 |
| $FeSO_4 \cdot 7\,H_2O$ | 5 | $Na_2MoO_4 \cdot 2\,H_2O$ | 0.2 |

### S5 Microwell Cultivations

Prior to experiments, all strains were stored at −80 °C in 10 $\%_v$ glycerol and plated on LA plates supplemented with antibiotics whenever applicable (25 $mg\,L^{-1}$ chloramphenicol for mScarlet-strains, 15 $mg\,L^{-1}$ gentamicin for sfGFP-strains, and 30 $mg\,L^{-1}$ kanamycin for non-fluorescent knock-out strains). Precultivations and actual cultivation experiments were performed in the mineral salt medium described in Table S6. No antibiotics were used in precultivations and experiments.

Figures S5 and S6 show the optical densities at 600 nm ($OD_{600}$) and fluorescence intensities corresponding to sfGFP and mScarlet emission maxima in the initial small-scale experiments, which were used to verify the growth of the double knock-out *A. baylyi* ADP1 Δ*gntT*:*E. coli* Δ*ptsI* consortium. All consortia accumulated $OD_{600}$ indicating growth, and the fluorescence intensities were considerable only when respective fluorescent protein genes were present in the culture. Methods related to these experiments are given in the main text.

### S6 Bioreactor Cultivations

#### S6.1 Results

To experimentally support the proposed carbon flow and to demonstrate the scalability of the carbon cross-feeding system, a bioreactor cultivation of Akr:Ekr (AbΔ*gntT*:EcΔ*ptsI* with mScarlet and sfGFP) was carried out with approximately hourly sampling. Figure S7 shows the measured optical densities, pHs and concentrations of glucose and acetate. For reference, cultivations with Eg as well as Akr:Eg and Ar:Eg were also carried out.

Akr:Ekg was capable growing in the bioreactor, demonstrating the scalability of the

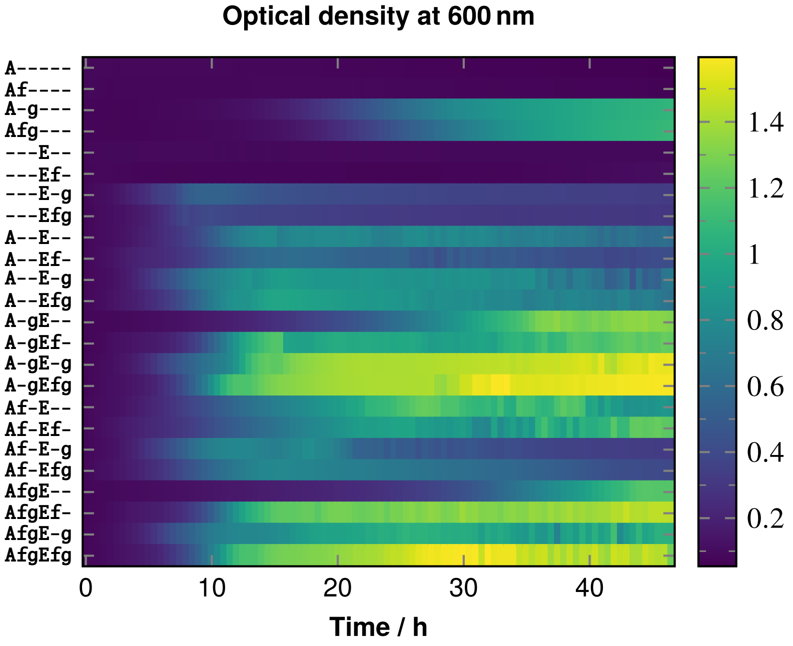

Figure S5: Optical density at 600 nm in the initial small-scale experiments. A denotes *A. baylyi* ADP1 and E *E. coli*. Fluorescent protein gene is denoted with f and the ability to utilize glucose with g. In effect, the wild-type *A. baylyi* ADP1 is denoted with A-g, and the mScarlet-carrying knock-out strain with Af-. Each horizontal lane corresponds to mean of three biological replicates cultivated in defined medium (Table S6) with 50 mM glucose at 30 °C.

proposed carbon cross-feeding. However, like in the small-scale cultivations, its growth rate was much lower than those of the other cultures. Akr:Ekg cultivation received an almost 5 mM residual acetate concentration from precultivations, which was consumed rapidly within the first 4 h. After this initial stage of growth, Akr:Ekg ceased to grow for approximately 6 h. Given that the wild-type *E. coli* accumulated acetate readily when cultured alone, it seems fair enough to assume that growth was restored due to acetate formed by Ekg. Comparing to the ODE framework and the numerical solutions obtained (Section S2), it seems likely that Akr:Ekg used most of the first 10 h on adapting its *A. baylyi* ADP1 to *E. coli* -ratio towards the optimum. Regardless of low growth rate, glucose was consumed by Akr:Ekg during the whole cultivation time. As the used *E. coli* strain Ekg was unable to utilize glucose, the drop in glucose concentration was due to glucose oxidation to gluconate by *A. baylyi* ADP1 Δ*gntT*. The steady descent of pH along with the steady drop of glucose concentration also supports gluconate production. Altogether, these experimental observations were consistent with the smaller-scale experiments, the ODE framework, and the proposed carbon flow (glucose to gluconate by *A. baylyi* ADP1, gluconate utilized and acetate excreted by *E. coli*, and acetate utilized by *A. baylyi* ADP1).

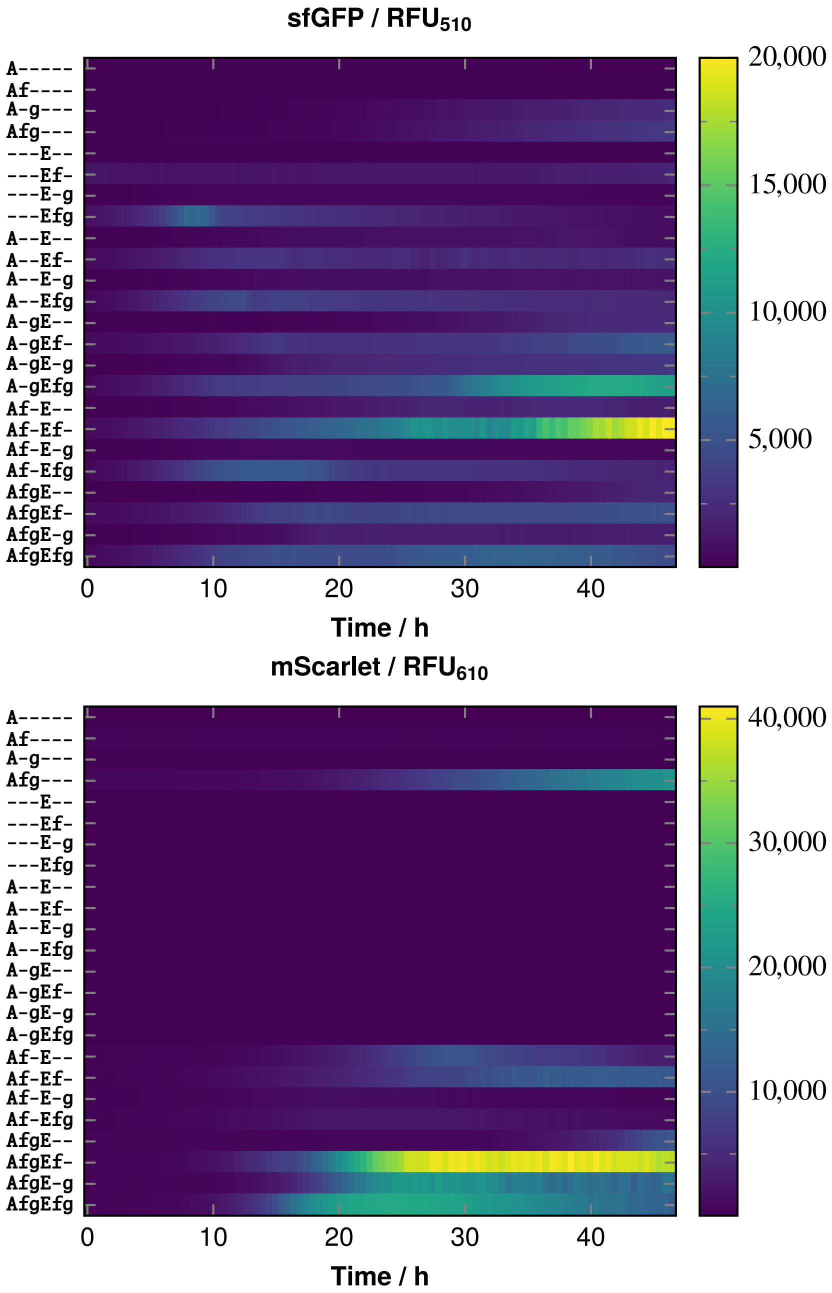

Figure S6: Fluorescence intensities at 510 nm and 610 nm in the initial small-scale experiments. A denotes *A. baylyi* ADP1 and E *E. coli*. Fluorescent protein gene is denoted with f and the ability to utilize glucose with g. For example the mScarlet-carrying knock-out *A. baylyi* ADP1 strain is denoted Af-. RFUs stand for the relative fluoresence units of the microwell plate reader. Each horizontal lane represents the mean of three biological replicates grown in defined medium (Table S6) with 50 mM glucose at 30 °C.

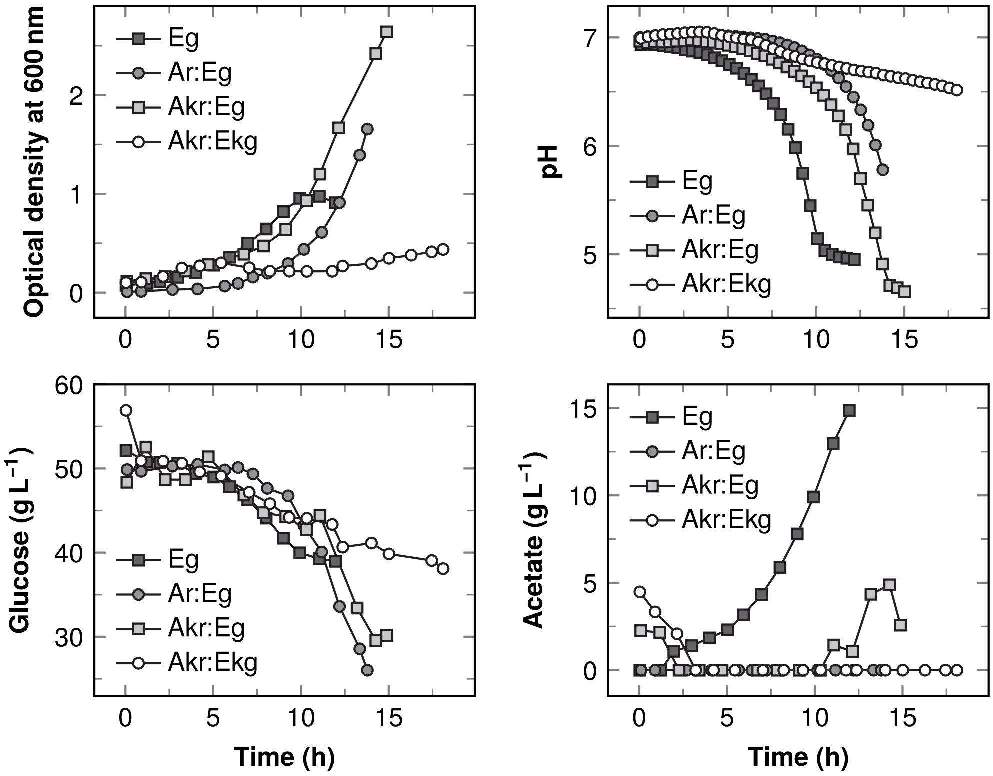

Figure S7: Bioreactor cultivations of the consortia and wild-type *E. coli*. Ar stands for *A. baylyi* ADP1 with mScarlet, Akr for ArΔ*gntT*, Eg for *E. coli* with sfGFP, and Ekg for EgΔ*ptsI*. Samples were drawn at approximately 1 h intervals. Neither biological nor technical replicates were made. $OD_{600}$ was measured with a spectrophotometer, glucose and acetate concentration were measured with a high-performance liquid chromatograph, and pH was continuously monitored by the reactor's control tower.

As for Eg and Akr:Eg, acetate accumulated towards the end of the cultivations, like in [21]. Eg accumulated acetate much earlier, which demonstrates the efficiency of acetate removal by Akr. Therefore it seems likely that acetate was the limiting factor in Akr:Ekg. Analogous to [22], increasing acetate excretion by *E. coli* might increase growth of the Akr:Ekg consortium. On the other hand, this also provided experimental support for not assuming a constant acetate concentration in *A. baylyi* ADP1 Δ*gntT*:*E. coli* consortia. Section S2 along with these findings indicate, that the two-way coupled *A. baylyi* ADP1 Δ*gntT*:*E. coli* Δ*ptsI* consortium should be more stable than the others. As a final note it is observed that culture acidity rose the quickest in Eg, but also markedly in Akr:Eg and Ar:Eg, which provides an explanation for steep descents of sfGFP fluoresence in some of the small-scale cultivations presented in main text (Figures 2 and 3).

#### S6.2 Precultivations

Precultivations for the reactor experiments were carried out in two stages. The first stage was started 24–36 h prior to reactor start-up as a 5 mL tube culture inoculated from LA-plates. In the second stage 4 mL of the tube-culture was used to inoculate 50 mL cultures in flasks 12–18 h prior to reactor inoculation. Both precultivation stages were incubated with 250 RPM or 300 RPM shaking at temperature of either 37 °C (*E. coli*) or 30 °C (*A. baylyi* ADP1). Precultures were performed with the same defined medium (Table S6) as the actual experiments. The wild-type *A. baylyi* ADP1 and *E. coli* strains were precultivated with 10 g $L^{-1}$ glucose. The knock-out *A. baylyi* ADP1 Δ*gntT* was precultivated with 2.5 g $L^{-1}$ glucose and 75 mM acetate, and *E. coli* Δ*ptsI* was precultivated with 50 mM gluconate. Glucose was used along with acetate in precultivating *A. baylyi* ADP1 Δ*gntT*, because its oxidation to gluconate is beneficial even if the gluconate is not utilized [23].

#### S6.3 Bioreactor configuration and operation

The reactor was a 1 L UniVessel Glass Culture Vessel (Sartorius, Germany) connected to a Biostat B plus control tower (Sartorius) monitoring temperature and pH. The reactor's impeller was a six-blade Rushton turbine with a diameter of 4.5 cm. A stirring rate of 350 RPM was used in all cultivations.

The targeted working volume was 0.55 L, but some variation occurred because of reactor sterilization in autoclave. Inoculum volumes of 50 mL were used such that the initial optical density would be at most 0.1. Equal amounts of strains in terms of optical density were used in inoculating consortia. The input rate of filtered air was set to 0.55 L $min^{-1}$ (gas at 20 °C temperature and 1.2 bar absolute pressure), correspoding to a volume-specific flow-rate of approximately 1 vvm. Temperature was maintained at 30 °C.

#### S6.4 Sample analyses

The reactor cultivations were sampled approximately hourly. 2 mL samples were drawn and divided to 1 mL aliquots. Right after sampling, optical densities at 600 nm and 700 nm were measured from one aliquot with a Ultrospec 500 pro spectrophotometer (Amersham Biosciences, UK). Because mScarlet has absorption maximum at 569 nm, it could bias the $OD_{600}$ measurements [24]. Therefore optical densities were measured at 700 nm as well. However, the bias at 600 nm remained neglicible.

The cells were removed from the other 1 mL aliquot by centrifugation, and concentrations of glucose and acetate were quantified from supernatants (stored at −20 °C and thawed) with a high-performance liquid chromatograph (HPLC). A LC-20 AD Prominence liquid chromatograph, SIL-20 AC Prominence auto sampler, and RID-10

A refractive index detector (Shimadzu, Japan) were used with a 30 cm Rezex RHM-Monosaccharide $H^+$ (8 %) column (Phenomenex, USA). The column was operated with 40 °C temperature and 0.6 mL min$^{-1}$ flow. Filtered (0.2 µm) 5 mM $H_2SO_4$ was used as the mobile phase.
